# Biomolecular phase boundaries are described by a solubility product that accounts for variable stoichiometry and soluble oligomers

**DOI:** 10.1101/2025.08.27.672390

**Authors:** Sk Ashif Akram, Aniruddha Chattaraj, Terry Salava, Jonathon A. Ditlev, Leslie M. Loew, Jeremy D. Schmit

## Abstract

The solubility product is a rigorous description of the phase boundary for salt precipitation and has previously been shown to qualitatively describe the condensation of biomolecules. Here we present a derivation of the solubility product showing that the solubility product is also a robust description of biomolecule phase boundaries if care is taken to account for soluble oligomers and variable composition within the dense phase. Our calculation describes equilibrium between unbound monomers, the dense phase, and an ensemble of oligomer complexes with significant finite-size contributions to their free energy. The biomolecule phase boundary very nearly resembles the power law predicted by the solubility product when plotted as a function of the monomer concentrations. However, this simple form is concealed by the presence of oligomers in the dilute phase. Accounting for the oligomer ensemble introduces complexities to the power law phase boundary including re-entrant behavior and large shifts for stoichiometrically matched molecules. We show that allowing variable stoichiometry in the dense phase expands the two phase region, which appears as curvature of the phase boundary on a double-logarithmic plot. Furthermore, this curvature can be used to predict variations in the dense phase composition at different points along the phase boundary. Finally, we show how the solubility product power law can be identified in experiments by using dilute phase dissociation constants to account for the oligomer ensemble.

## Introduction

Biomolecular phase separation has emerged as an important mechanism of cellular organization.^1,2^ In some cases biomolecular condensate formation is driven by the phase separation of a single dominant scaffold molecule that occurs at a threshold concentration *C*_*sat*_.^3–6^ However, in many other cases the formation of biomolecular condensates requires multiple scaffold molecules and the condensation threshold depends on the concentration of all of the scaffolds. ^7–16^ A common case is when two scaffold species are required, which is analogous to salt precipitation where both anion and cation components are required. In contrast to the *C*_*sat*_ describing single-component phase separation, salts remain soluble when the product of the ion concentrations remains below a threshold value *K*_*SP*_, known as the solubility product constant. Application of the solubility product concept to biological condensates has been promising,^16–19^ however, factors like conformational diversity and high valency introduce complexities not present in simple salts that have confounded interpretation of experiments. For example, phase separating biomolecules often form oligomers that remain in the dilute phase.^19,20^ Additionally, it can be difficult to determine the effective valence of disordered proteins. Even in cases where the valence can be obtained from well-defined interaction modules, asymmetries due to non-interacting “spacers”^21^ or electrostatic interactions can result in unexpected mixing ratios. Finally, biomolecular condensates tend to form without organized, repeating structural elements, which contributes to their liquid properties. This allows for heterogeneity in stabilizing interactions and variation in the stoichiometric composition of the condensed phase. This is biologically significant because stoichiometric variation has been linked to the functional state of condensates. ^22–25^

Here we present a theory of the solubility product that accounts for many of the complexities inherent to biomolecular phase separation, specifically soluble oligomers and variable stoichiometry in the dense phase. We show that the phase boundary is most naturally defined in terms of the concentrations of free monomers. The presence of oligomers in the dilute phase complicates measurement of the monomer concentrations and leads to phenomena like the “magic number” effect^26–29^ and re-entrant transitions.^30–33^ Fluctuations in the dense phase composition expand the two-phase region of the phase diagram, which appears as curvature in the phase boundary when viewed on a double-logarithmic scale. This relationship can be inverted such that curvature in the phase boundary can be used to predict how composition of the dense phase changes with the solution composition. Finally, we show that the optimal stoichiometric composition of the condensed phase requires a balance of chemical potentials that does not coincide with balanced valences in the dilute phase (except in the special case of 1:1 stoichiometry). Together these results show how measurement of the dilute phase can be informative about the properties of the condensed phase.

## Results

### Model describes the co-assembly of two molecular species

We present theory and simulations for a solution consisting of two macromolecular species, that we refer to as “A” and “B”, immersed in a solvent that is implicitly accounted for by the interaction energies. The molecules can assemble into complexes that are each characterized by a free energy *F* (*n, m*) where the argument *n* indicates the number of A molecules and *m* is the number of B molecules in the cluster. We use the term “cluster” generically to indicate a reversible, in the thermodynamic sense, supramolecular complex without implications of the solid/liquid or oligomer/phase properties of that complex.

The grand partition function of the solution is

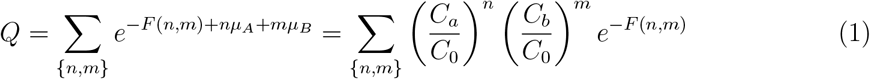

Where the second step takes the monomers to be the reference state (i.e., *F* (1, 0) = *F* (0, 1) = 0) and assumes the solution to be sufficiently dilute that the chemical potentials are *µ*_*A*_ = ln(*C*_*a*_*/C*_0_) and *µ*_*B*_ = ln(*C*_*b*_*/C*_0_), where *C*_0_ is a reference concentration that we take to be 1 M in our comparison to simulations. The chemical potentials, free energies of association *F* (*n, m*), and all subsequent energies, are expressed in units of *k*_*B*_*T*, where *k*_*B*_ is Boltzmann’s constant and *T* is the absolute temperature. Hereafter, we work in terms of dimensionless concentrations denoted with a lowercase *c*, as in *c*_*a*_ = *C*_*a*_*/C*_0_, *c*_*b*_ = *C*_*b*_*/C*_0_, etc. The ideal treatment of the chemical potentials is justified by the fact that monomers, by definition, are not interacting with other molecules. However, in crowded environments, such as the cytoplasm, the concentration rescaling can absorb the activity coefficient of the dispersed phase. In our notation concentrations with a lowercase subscript indicate the monomer concentration while capital subscripts are reserved for the total concentration (with a superscript, if necessary, to indicate the dense or dilute phase).

#### Divergence in the partition function indicates the phase boundary

We divide the partition function into terms representing the monomers, oligomers, and condensed phase.

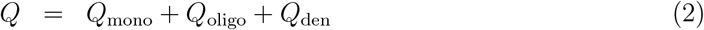

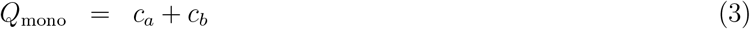

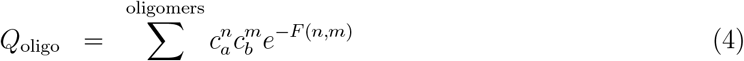

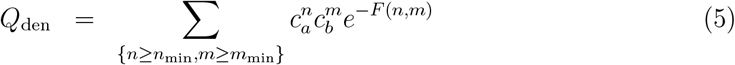

We consider a cluster to be part of the dense phase when it exceeds threshold sizes *n*_min_ and *m*_min_ that are large enough that surface corrections to the free energy can be neglected. In other words, for clusters greater than these cutoffs the free energy scales linearly with size (i.e., *F* (*λn, λm*) = *λF* (*n, m*) for *λ* ≥ 1, *n > n*_min_, and *m > m*_min_). All clusters that do not satisfy these conditions are included in *Q*_oligo_.

We first consider the case where the stoichiometry of the dense phase is tightly constrained, which is valid in the tight binding limit where all bonds are satisfied in the dense phase. In this case, the number of B molecules in the dense phase scales in direct proportion to the number of A molecules, *m* = *sn*, where *s* is the stoichiometric ratio. The Boltzmann factors above the size cutoff can be written as 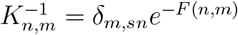, where *δ* is the Kronecker delta and 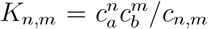 is the equilibrium dissociation constant for the (*n, m*) cluster.

Applying this constraint to *Q*_den_, the summation over *m* can be evaluated

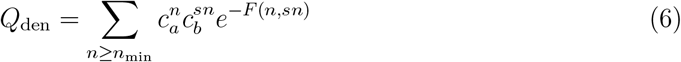

Next, we use the fact that the free energy scales linearly with size to write *F* (*n, sn*) ≃ *nf*_*A*_ where *f*_*A*_ = lim_*n*→∞_ *F* (*n, ns*)*/n* describes the free energy per A molecule in large clusters.

The summation is evaluated as

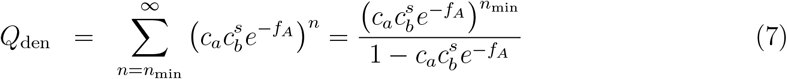

Note that *Q*_den_ diverges when *f*_*A*_ − ln *c*_*a*_ − *s* ln *c*_*b*_ = 0, which can be rearranged in the form of a solubility product

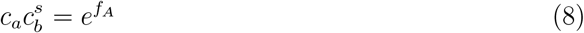

The requirement that the partition function is finite and positive means that the concentration product 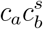 must be less than 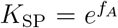. This limit on the monomer concentrations is the phase boundary. For a predetermined value of *c*_*a*_, the critical concentration of B molecules is *c*_*b*_ = (*K*_SP_*/c*_*a*_)^1*/s*^ which is a straight line with slope −*s*^−1^ on a double logarithmic plot. Note that omitting the small *n, m* terms from the *Q*_den_ sum does not affect condition for the phase boundary because *n*_min_ does not appear in Eq. 8.

### Oligomers perturb the phase boundary by suppressing monomer concentrations

In many experiments the known quantities are the total concentrations and the partitioning between the dense and dilute phases. In such experiments the phase boundary is plotted as a function of the total concentrations *c*_*A*_ and *c*_*B*_. Such phase boundaries do not show the expected power law behavior because the soluble fraction includes both monomers and oligomers.^34^ Soluble oligomers can cause qualitative differences between the apparent phase boundary and the solubility product in both the subsaturated regime and superstoichiometric mixtures.

To address these cases we note that, for suitably large *n*_min_, *Q*_den_ behaves like a step function because values of 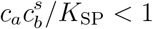 are strongly suppressed by the exponent *n*_min_ in the numerator of Eq. 7 (see Fig. 1A). Therefore, below the solubility threshold it is reasonable to approximate *Q ∼ Q*_mono_+*Q*_oligo_. Furthermore, in many cases *Q*_oligo_ will be dominated by a small number of favorable species. ^18,35^ Therefore, *Q*_oligo_ can be approximated by truncating the cluster summation to include only the most significant oligomers for a given system. The selection of oligomers to include will be highly system dependent. In some cases the range of allowable oligomers is determined by the molecular valences, as we demonstrate below in our comparison to experiments. In other cases, the strong dependencies arising from the exponential Boltzmann weights and concentration power laws will create a peak in the distribution of oligomers.

**Figure 1.**
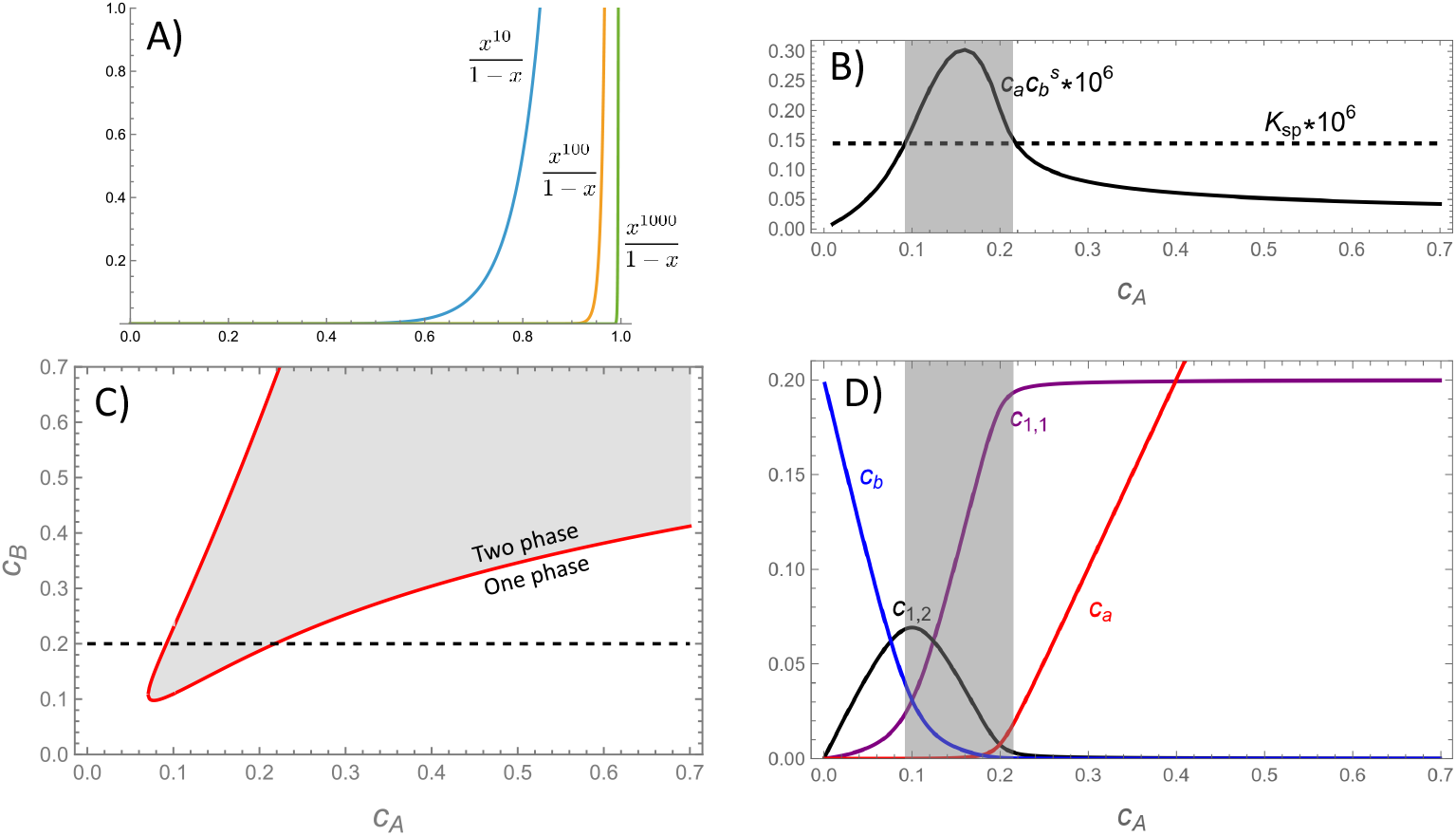
(A) The dense phase partition function *Q*_den_ has the characteristics of a step function, which allows the approximation *Q* ≃ *Q*_mono_ + *Q*_oligo_ in the sub-saturated regime. Larger values of *n*_min_ increase the steepness of the step, while smaller values lead to more gradual phase transitions. (B) Plot of the monomer product 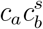 for fixed *c*_*B*_ (dotted line in panel C) computed from Eqs. 13 and 12. The product exceeds the threshold for condensation in the shaded region. (C) The phase boundary can be computed from the values where the solubility product reaches the *K*_SP_ threshold. (D) Plots of the oligomer concentrations reveal that the re-entrant behavior seen in panel C is due to an excess of 1:1 or 1:2 oligomers. Note that the oligomer concentrations in the shaded regions are not reliable because this calculation does not account for molecules in the dense phase. *s* = 1.4, *K*_1,1_ = *e*^−15*/s*^, *K*_2,1_ = *e*^−15^, and *f*_*A*_ = −15.75.

Once the approximate form of the partition function is obtained, it will be a polynomial that can be solved for *c*_*a*_ and *c*_*b*_. This solution can be used to solve for the locus of points where 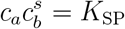, which corresponds to the phase boundary. To illustrate this approach, we consider a case *s* = 1.4 that, in the strong binding limit, corresponds to molecular valences that are related by *v*_*A*_ = *sv*_*B*_. When the A molecule is present in excess, we expect the dominant oligomer will be the 1:1 heterodimer, which fully saturates the B binding sites. Under conditions of excess B the dominant oligomer will be the 1:2 heterotrimer, which fully saturates the A binding sites. Thus, the partition function in the sub-saturated regime (i.e., no dense phase) can be approximated as

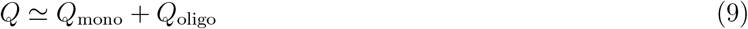

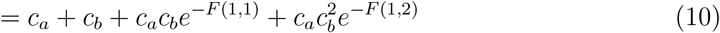

The total concentration of B molecules is given by

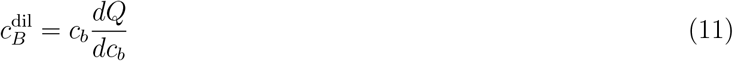

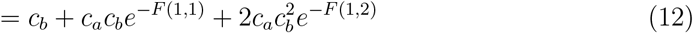

Similarly, the concentration of A monomers *c*_*a*_ is related to the total A concentration 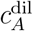 by

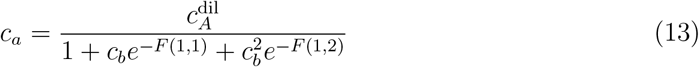

where the denominator is the binding polynomial for A molecules.^36^ Plugging Eq. 13 into Eq. 12 gives a cubic equation for *c*_*b*_. The real, positive root of this equation gives *c*_*b*_ as a function of 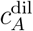 and 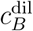, which can be used with Eq. 13 to give *c*_*a*_. The existence of a real, positive root is guaranteed by the positive coefficients in the *Q*_oligo_ sum, regardless of the *s* value or the particular oligomers included in the model.

Fig. 1B shows the solubility product 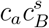 obtained from Eqs. 12 and 13 as a function of *c*_*A*_ for a fixed value of *c*_*B*_ (indicated by the dotted line in Fig. 1C). At low and high concentrations of A the solubility product lies below the solubility threshold *K*_SP_, indicating a single phase system. However, at intermediate concentrations, indicated by the shaded region, the solubility product rises above *K*_SP_. The locus of points where 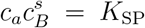 indicates the phase boundary, which is plotted as a red line in Fig. 1C. Fig. 1D shows how the population of oligomers shifts along the path shown in panels B and C (with the caveat that accounting for the dense phase will reduce the concentration in the shaded, two-phase region). In agreement with the assumptions of the model, the low and high concentration regimes are dominated by oligomers where the valence of the minority component is saturated. This saturation is responsible for the re-entrant behavior observed in Fig. 1C.

#### Oligomer formation maximizes the translational entropy of stoichiometry imbalanced solutions

The re-entrant behavior can be understood as follows. On the left edge of Fig. 1C, where the B:A ratio is very large, the system optimizes its free energy by maximizing the translational entropy while saturating the A binding sites with 1:2 heterotrimers. In the large *c*_*b*_ limit, the abundance of B molecules means that the translational entropy penalty for recruiting B molecules to a cluster is smaller than the benefit of forming additional bonds. This drives *c*_*a*_ to very low values such that 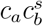 falls below the *K*_SP_ threshold. This can be seen in the theory by noting that, for large *c*_*b*_, the 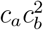 scaling of the trimer dominates over the 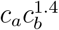 scaling of the dense phase.

More generally, for systems with saturating bonds, there will always be an oligomer where the 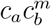 where the A bonds are saturated and the *m* exponent is greater than the 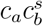 exponent of the dense phase. This is because contiguity of a dense phase requires that B molecules are shared between nearby A molecules. However, not all systems have saturating bonds. For example, “scaffold-client” architectures^37^ (also called “hub-driver” ^38^) are capable of forming homotypic condensates that do not disperse in the presence of excess driver.^24,39^

When the concentration of A molecules increases such that the B:A ratio is comparable to the valence ratio, the dilute phase becomes populated by oligomers close to the stoichiometry of the dense phase. In the case of our *s* = 1.4 toy model, the dominant oligomer is the 1:1 heterodimer. In these conditions, the free monomer concentration is set by the dissociation constant of the most favorable oligomer. This has two implications. First the dissociation constant describes the concentration at which the translational entropy cost of forming the oligomer balances the bonding energy gains. Therefore, a partial set of bonds, which occurs with the 1:1→1:2 transition, will not be sufficient to overcome the translational entropy cost. Second, for a system to form a condensate, *K*_SP_ must be lower than the dissociation constant of the comparable oligomer *s* ≃ *m/n* due to the additional bonds and/or bonding entropy available in the dense phase. Therefore, the 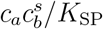 term of the dense phase dominates over the 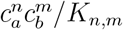 weight of the oligomer.

Finally, when the system reaches *c*_*a*_ ≫ *c*_*b*_, we arrive at the mirror image of the *c*_*b*_ ≫ *c*_*a*_ case, where the dilute phase fully satisfies the bonds of the B molecules while maximizing the translational entropy. Here *c*_*b*_ ≃ 0, so the *c*_*a*_*c*_*b*_ weight of the dimer dominates over the 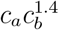 weight of the dense phase (which is further hindered by the 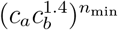 surface penalty).

### Stoichiometry matched oligomers increase sharply in concentration

In general, we expect that the molecular affinity in the dense phase, *f*_*A*_, will be more attractive than the per molecule oligomer affinities *F* (*n, m*)*/n* due to the additional conformational entropy and binding combinatorics when each molecule is surrounded by interaction partners. However, the dense phase is inhibited by the large translational entropy penalty coming from the 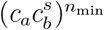 factor in the numerator of *Q*_den_. Therefore, condensation begins when the affinity advantage (*f*_*A*_ vs. *F* (*n, m*)) overwhelms the mixing entropy penalty, at which point *Q*_den_ dominates *Q*_oligo_. However, the transition between oligomer accumulation and dense phase formation is highly sensitive to the molecular details. In particular, previous work has shown that large shifts in the phase boundary can occur when the valences of the two species differ by an integer ratio, which has been called the “magic number effect”. ^26–29^

To understand the large shift in the phase boundary at integer valence ratios, we take the simplest case where the A and B molecules have nearly matched valences (the general case is shown in the Appendix). For closely matched valences we can write *s* = 1 + ϵ. Using *E* as a small parameter, the phase boundary, Eq. 8, can be written as 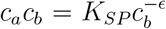. The dominant oligomer species will be the heterodimer, the concentration of which is given by *c*_1,1_ = *c*_*a*_*c*_*b*_*/K*_1,1_. Therefore, the concentration of dimer at the phase boundary is

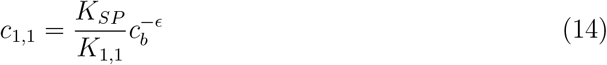

Phase separation will occur most readily when the dimer concentration is low, because this leaves more monomers to reach the *K*_*SP*_ threshold. In agreement with this expectation, we see that highly attractive dense phases (small *K*_*SP*_) lower the dimer concentration, while high affinity dimers (small *K*_1,1_) increase it.

Consider the case where the A molecules are unchanged and ϵ = *s* − 1 is varied by changing the B valence. We further assume that the solution contains an equimolar mixture of molecules such that *c*_*a*_ = *c*_*b*_ = *c*_*ab*_. This allows us to rewrite the threshold dimer concentration as

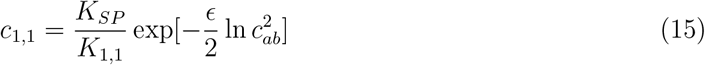

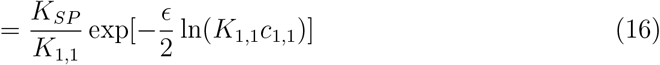

Next, we recursively plug Eq. 16 back into itself to obtain

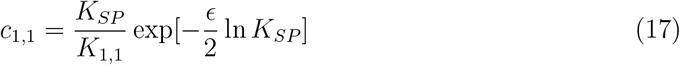

where we have dropped terms of order ϵ ^2^. While it is possible to sum the perturbation series to obtain an exact solution (see Supporting Information), the first approximation (Eq. 17) is sufficient to see the magic number phenomenon. Eq. 17 can also be written in terms of the cluster free energies

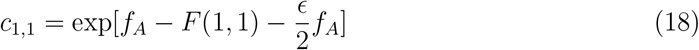

In the tight binding limit, the dimer free energy is determined by the maximum number of bonds that can be formed. Therefore, *F* (1, 1) = *f*_1_ min(*v*_*A*_, *v*_*B*_) = *f*_1_*v*_*A*_ min(1, (1 + *E*)^−1^), where *f*_1_ is the energy of a monovalent bond. Therefore, for *E*< 0 the threshold dimer concentration varies as

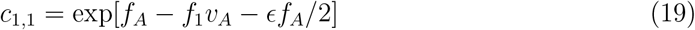

which increases exponentially as ϵ approaches zero. For ϵ *>* 0 we have *F* (1, 1) = *f*_1_*v*_*A*_/*s* ≃ *f*_1_*v*_*A*_(1 − ϵ), so the threshold dimer concentration varies as

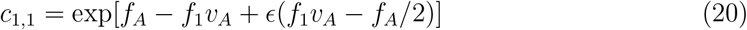

which *decreases* exponentially provided that *f*_1_*v*_*A*_ *< f*_*A*_/2. This means that *c*_1,1_ has a cusp at *E* = 0 with exponential drops on either side (Fig. 2A). The large increase dimer concentration depletes the monomer pool, which causes the phase boundary to shift to much higher concentrations (Fig. 2B).

**Figure 2.**
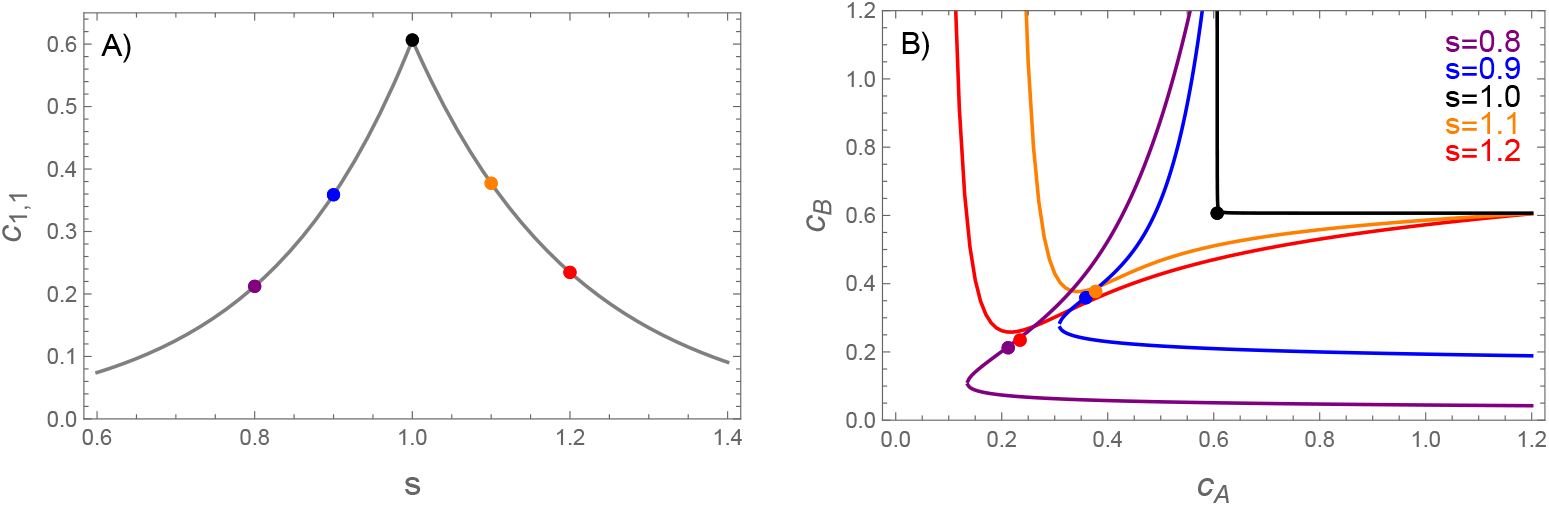
A) The heterodimer concentration at the phase boundary (computed from Eqs. 19 and 20) has a cusp peak at *s* = 1. B) As a result of this peak, the phase boundary shows a large shift at *s* = 1. The colored dots show the dimer concentrations at the points labeled in panel A. The dots lie very close to the phase boundary, indicating dimers account for most of the molecules in the dilute phase. *F* (1, 1) = *f*_1_*v*_*A*_ min(1, 1/*s*), *f*_*A*_ = 1.05*v*_*A*_*f*_1_, *v*_*A*_*f*_1_ = 10

Inspection of Eqs. 19 and 20 allows us to extend these arguments outside of the tight binding limit. The dense phase translational entropy, represented by the *f*_*A*_/2 term, will always favor larger B valences. Similarly, the oligomer free energy *F* (1, 1) will also favor large *v*_*B*_, but with substantially diminishing returns once the B valence exceeds *v*_*A*_. Therefore, we can expect a magic number peak in the phase boundary whenever the change in the slope of *F* (1, 1) is large compared to *f*_*A*_/2. This will not be true in the weak binding regime where entropic contributions dominate the change in *F* (1, 1) or in cases where molecular geometry restricts bond saturation in the dimer. ^27,35,40^ However, in such cases there may be a larger oligomer that has a magic number peak at the phase boundary.^35^ In the Supporting Information we derive an exact expression for the concentration of an arbitrary oligomer at the phase boundary

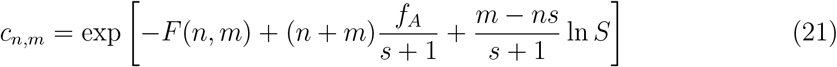

where *S* = *c*_*b*_/*c*_*a*_ is the ratio of monomers at a particular point along the phase boundary. The first two terms of Eq. 21 reflect a competition between the oligomer free energy and the dense phase free energy. While the dense phase is generally more favorable, for small values of *n* and *m* the difference can be small enough that oligomers are significantly populated. Large oligomers, however, are exponentially suppressed until their size becomes large enough that *F* (*n, m*) becomes comparable to the dense phase free energy. Therefore, while Eq. 21 describes magic number peaks for all oligomers, in practice only oligomers with integer stoichiometric ratios are present at concentrations high enough to significantly perturb the phase boundary (see SI).

### Solubility with soft stoichiometry constraint

Thus far, we have focused on the strong binding limit where the stoichiometry of the dense phase, and oligomers to a lesser extent, is restricted by the prohibitive cost of unsatisfied bonds. However, biomolecules typically are conformationally flexible, form bonds with affinities on the order of *k*_*B*_*T*, and form condensates with liquid-like disorder. Therefore, unlike salts, biomolecular condensates are not restricted to a specific stoichiometry. To understand the effects of compositional variation in the dense phase, we begin with the dense phase partition function, Eq. 5. The summations in Eq. 5 encompass the upper right quadrant *n, m >* 0, which can also be covered using polar coordinates

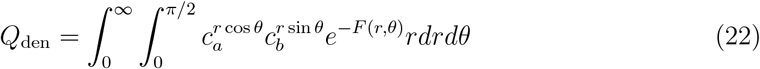

The line of ideal stoichiometry *m* = *sn* can be expressed in terms of an angle tan *θ*_0_ = *s* (see Fig. 3A). We expect that the free energy *F* (*r, θ*) will restrict the significant contributions to the integral to regions near the ideal stoichiometry. Therefore, we expand the condensed phase free energy to lowest order in *δ* = *θ* − *θ*_0_.

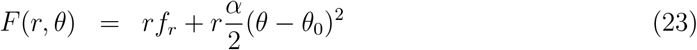

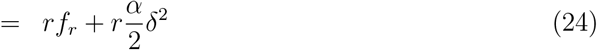

**Figure 3.**
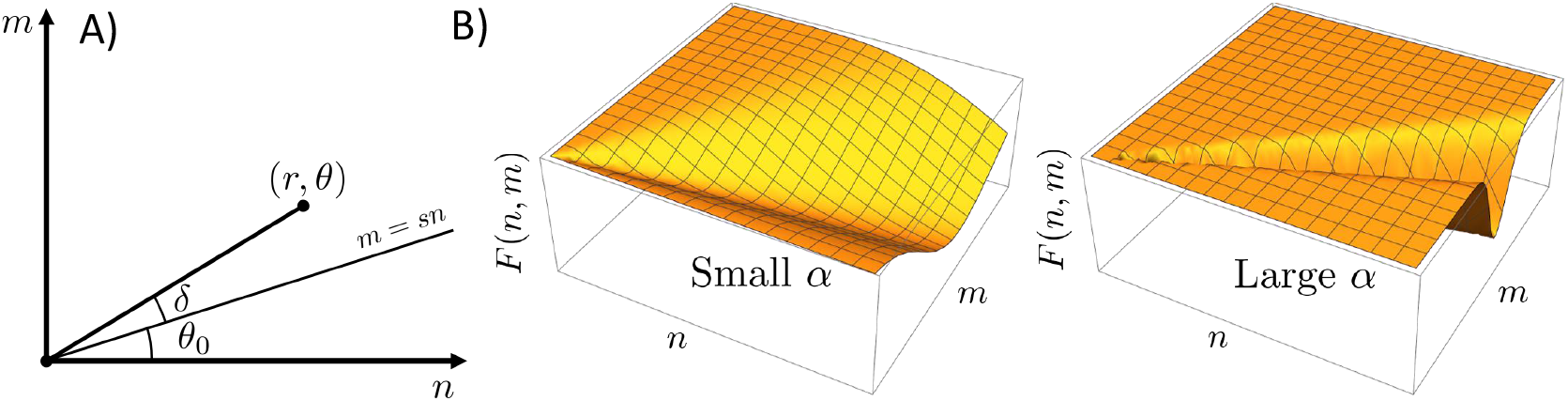
(A) Coordinate change from cartesian axes (*n, m*) to polar coordinates (*r, θ*). The radial integration is performed over the angle *δ* = *θ* − *θ*_0_, which is measured relative to the line of optimal stoichiometry. (B) Schematic plots of *F* (*r, θ*) for small and large values of *α*. Larger values of *α* correspond to narrower low energy troughs in the free energy landscape and clusters that are more resistant to changes in composition.

Here *f*_*r*_ describes how the free energy scales with *r* when the condensate has the ideal stoichiometry

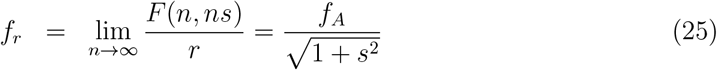

and *α* is a parameter that describes how the free energy varies as the stoichiometry deviates from the ideal ratio

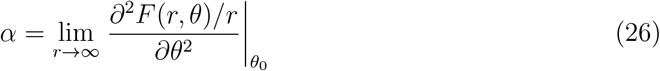

The tight stoichiometry limit can be recovered in the limit *α* → ∞. The gaussian approximation in Eq. 24 will be valid when the stoichiometry fluctuations are small enough that anharmonic terms (*O*(*δ*^3^)) in the free energy are negligible.

For consistency with the expansion of *F* (*r, θ*), we also Taylor expand the chemical potential terms to second order in *δ*. The partition function with the expanded free energy takes the form (see SI for details)

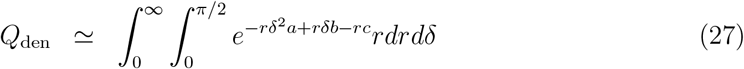

where

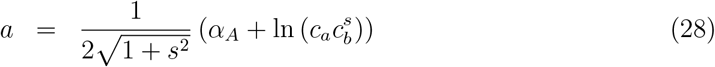

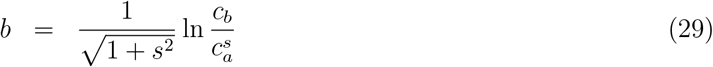

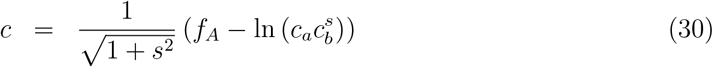

and 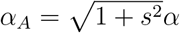. Evaluation of the integrals in Eq. 27 (see SI) yields

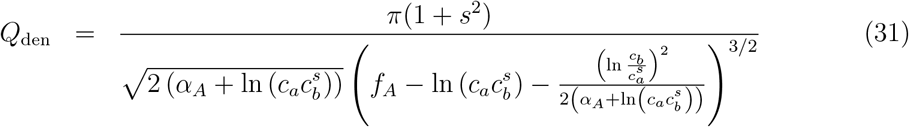

#### Composition fluctuations expand the two-phase regime

The phase boundary lies at the point were the partition function diverges. Inspection of Eq. 31 shows that this occurs when

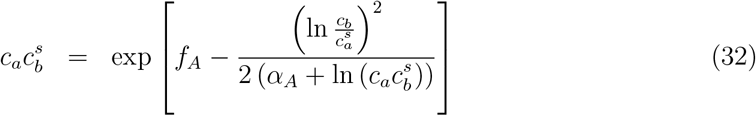

Using the recursive approximation ln 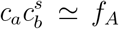, we can define the effective resistance to composition fluctuations *α* ^′^ = *α*_*A*_ + *f*_*A*_. This simplifies the solubility threshold to

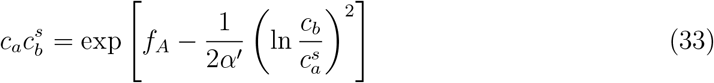

Eq. 33 has nearly the same form as the expression we found for the solubility product in the limit of a tight stoichiometry constraint (Eq. 8). The difference is a correction term that accounts for fluctuations in the composition of the dense phase. Note that the fluctuation term is always negative since *α* ^′^ must be positive for the gaussian integral to converge. Therefore, the fluctuation term lowers the free energy of the dense phase relative to the tight stoichiometry case. This is expected because loosening the stoichiometry constraint contributes additional terms to the partition sum, which must lower its free energy.

Eq. 33 can be solved for *c*_*b*_ (see SI), which yields an expression for the phase boundary

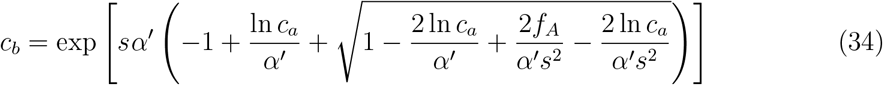

Eq. 34 is plotted for various values of *α* in Fig. 4. We see that decreasing *α* has the effect of introducing curvature in the phase boundary when plotted on double logarithmic axes.

**Figure 4.**
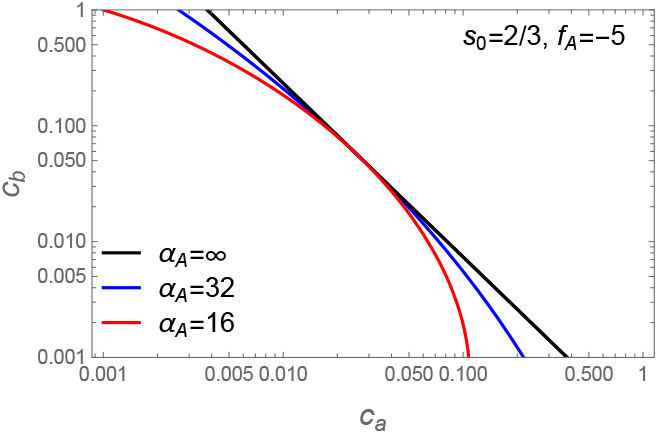
Plot of Eq. 34 for *s* = 2/3. The *α* = ∞ limit is linear with slope − *s*^−1^ on a log-log axes (black lines) due to the power law behavior of the tight stoichiometry case (Eq. 7). Lower values of *α* introduce curvature to the phase boundary and increase the phase separated region (upper right) at the expense of the single phase region (lower left). The curved phase boundary is tangent to the black line when *s* ln *c*_*a*_ − ln *c*_*b*_ = 0.

#### Phase boundary curvature describes variations in the dense phase stoichiometry

Our next task is to determine how the composition of the dense phase varies with changes in the solution stoichiometry. In cases where the concentrations of the dense phase are known, this calculation is equivalent to determining the tie lines connecting the dense and dilute phase.

The total concentration of dense phase molecules can be computed using

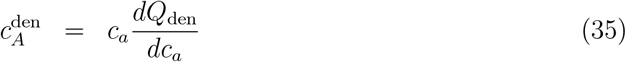

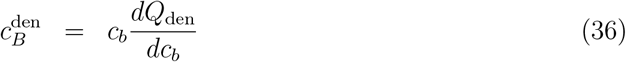

Evaluation of the derivatives gives the stoichiometric ratio (see SI)

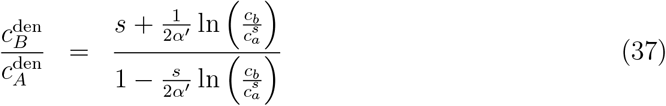

Both the numerator and denominator have correction terms that are proportional to *α* ^′ −1^, so they become negligible in the tight stoichiometry limit (*α* → ∞). In this limit we recover the optimal stoichiometry, *s*. Additionally, we obtain the optimal stoichiometry when *s* ln *c*_*a*_ − ln *c*_*b*_ = 0. This expression describes the balance of chemical potentials required for a growth increment of Δ*n* type A molecules to incur the same mixing entropy penalty as recruiting *s*Δ*n* molecules of type B. Notably, this condition (i.e.,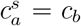) is different than the more intuitive expectation that the dense phase has the optimal stoichiometry when the dilute phase has a matching stoichiometry *sc*_*a*_ = *c*_*b*_ (except for the special case *s* = 1). When *s* ln *c*_*a*_ − ln *c*_*b*_ /= 0 the dense phase stoichiometry is perturbed such that the molecule that is present in excess in the dilute phase is also over-represented in the dense phase.

Comparison of Eqs. 33 and 37 reveals that a single parameter, *α* ^′^, controls both the compositional variation in the dense phase and the deviation of the phase boundary from a −*s*^−1^ power law. Therefore, the curvature of the phase boundary, when plotted on double logarithmic axes can be used to predict the composition of the dense phase.

### Simulations of sticker-spacer polymers show a curved phase boundary

To test the predicted relationship between the phase boundary and the dense phase composition, we conducted simulations of a bead-spring polymer model. Within each chain, certain beads behave as stickers while others serve as spacers to impart flexibility^21^ (Fig. 5A). Our model only includes heterotypic interactions in which a bond requires one sticker of type A and one of type B. The number of stickers per molecule defines the molecular valences, *v*_*A*_ and *v*_*B*_.

**Figure 5.**
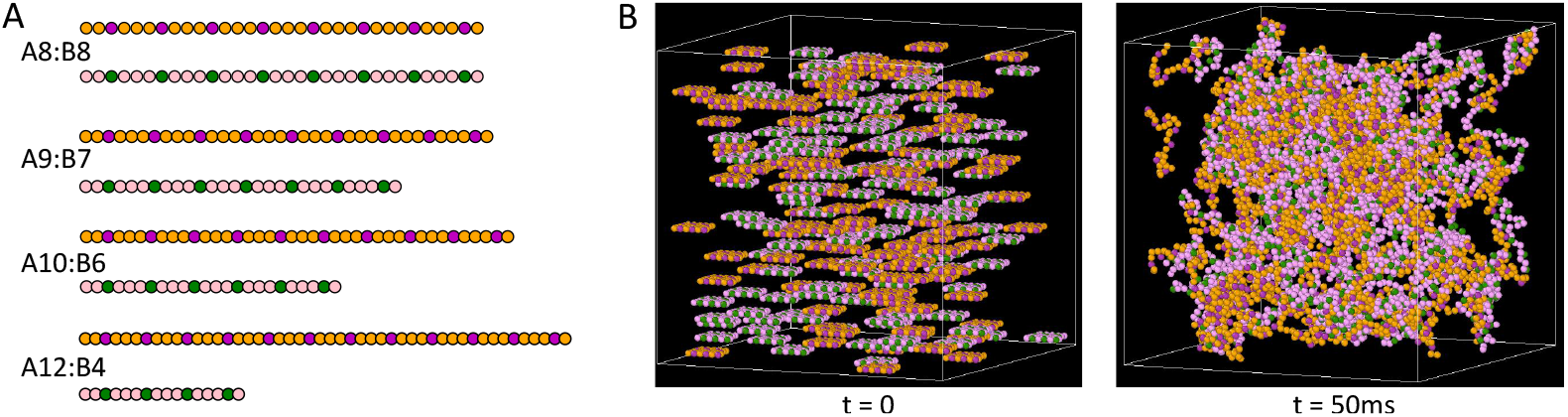
(A) Schematic representations of the bead-spring polymers used in the SpringSaLaD simulations. Green and magenta beads represent the A and B-type stickers, respectively, while the orange and pink beads represent spacer segments. (B) Snapshots of the A8:B8 system with *N*_*A*_ = *N*_*B*_ = 120 molecules at the start of the simulation and after 50ms.

Each simulation starts with a predetermined number of each molecule, uniformly distributed in a cubic simulation volume. The systems evolve by Langevin dynamics using the SpringSaLaD software package (see SI for details).^41^ As a result of binding interactions, these multivalent molecules form clusters, where a cluster includes all beads connected by intra-chain or inter-molecular bonds (Fig. 5B).

We first establish the convergence of our simulation by tracking the time evolution of the cluster size distribution and bond saturation (Fig. 6A-C). Multiple stochastic runs are performed to extract an average behavior of the system. Fig. 6A shows how the system approaches steady state, where the system bifurcates between a small fraction of molecules in very small clusters and a large fraction in clusters containing *>* 20 molecules (the total number of A+B molecules is 240). Figs. 6B and C display the average clustering and binding saturation dynamics. Once the system fluctuates around a steady state, we sample the cluster size distribution that consists of monomers (unbound molecules), oligomers (smaller clusters) and large clusters. Fig. 6D shows the extent of bond saturation as the clusters grow large in size showing that, while small clusters vary considerably in bond saturation, large clusters converge to a constant value of about 0.45.

**Figure 6.**
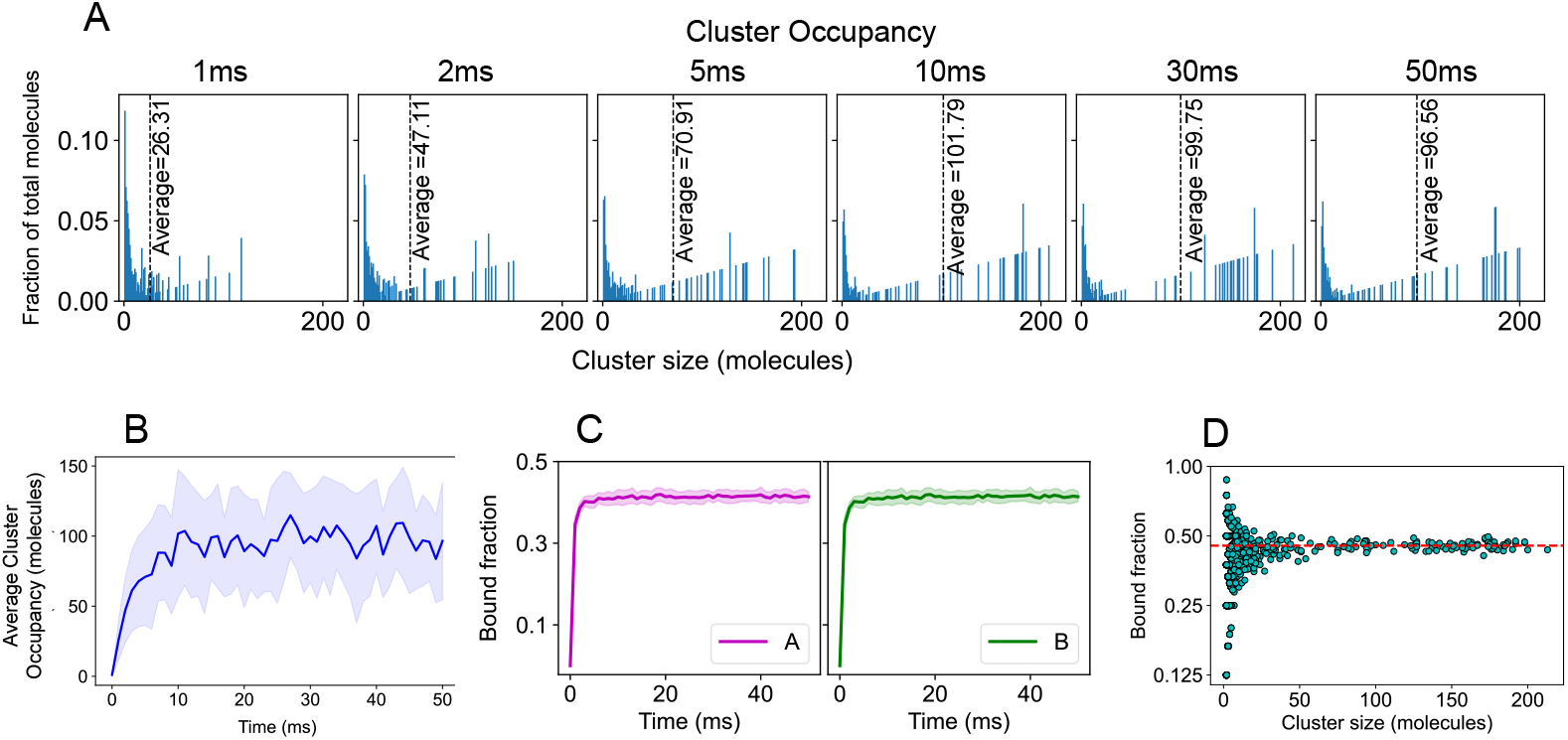
SpringSaLaD simulation of A8:B8 system. (A) Histograms of the normalized cluster occupancy, which we define as the fraction of molecules (*Nc*_*N*_ /(*c*_*A*_ + *c*_*B*_)) in clusters of size *N*, where *N* = *n* + *m* is the total number of A and B molecules in the cluster and *c*_*N*_ is the concentration of all clusters of that size *c*_*N*_ = ∑_*n*+*m*=*N*_ *c*_*n,m*_. The average cluster occupancy (dotted line) grows initially before converging to a constant value at about 10 ms. Each panel shows an average over 25 stochastic runs. (B) Time evolution of the average cluster occupancy averaged over 25 runs. The solid line shows the mean and the shaded area is standard deviation. (C) Bonded sticker fraction as a function of time, averaged over 25 runs. (D) Bonded sticker fraction as a function of cluster size obtained from 4 times (20, 30, 40, 50ms) across 25 runs (total 100 frames). Length of the cubic simulation box = 120 nm and molecular count, *N*_*A*_ = *N*_*B*_ = 120, molecular concentration ∼ 115 *µ*M.

To obtain the phase boundary, we varied the number of A and B molecules across eight values for a total of 64 mixtures. Referring to Eq. 8, we recall that the phase boundary is defined in terms of the monomer concentrations. Accordingly, Fig. 7A,C shows that the monomer concentrations are clustered along a curved line. The absence of points above the line indicates the presence of a phase boundary. The scatter of points below and/or left of the boundary is due to either initial conditions that are sub-saturated or highly supersaturated simulations where obtaining an accurate monomer concentration would require correcting for the volume occupied by the dense phase. To avoid these complications, we approximate the phase boundary by the points along the upper edge of the cluster.

**Figure 7.**
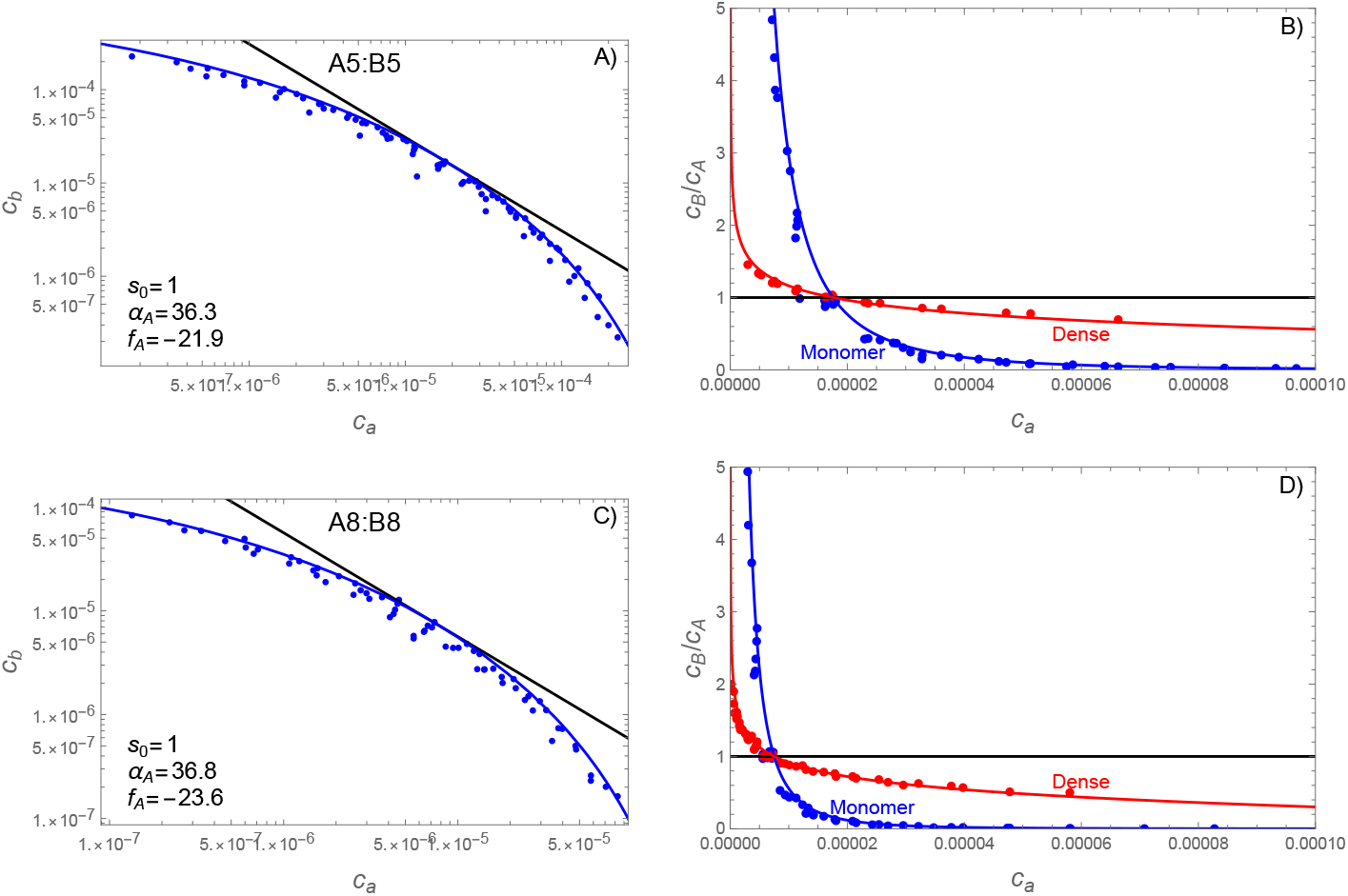
SpringSaLaD simulations reveal a curved phase boundary that predicts composition variations in the dense phase. Panels A,B show molecules with 5 stickers per chain (*v*_*A*_ = *v*_*B*_ = 5) while panels C,D show *v*_*A*_ = *v*_*B*_ = 8. Left panels (A,C) show the average concentration of free monomers for each of the simulated mixing ratios. Fitting Eq. 34 to the uppermost points yields values for *f*_*A*_ and *α*_*A*_. Right panels (B,D) show the stoichiometric ratios B:A for both the monomers (blue) and dense phase (red). Note that the monomer lines and data (blue) are equivalent between panels with only a change in the vertical axis. The monomer ratio is calculated from Eq. 34 divided by *c*_*a*_ and the dense phase ratio is calculated from Eq. 37 with Eq. 34 substituted for *c*_*b*_. The stoichiometric composition of the dense phase agrees well with the composition of clusters *n* + *m* ≥ 20 molecules observed in the simulations. In all panels simulation data are shown as points, theory as a solid line, and concentrations are given in the dimensionless units *c*_*i*_ = *C*_*i*_/(1 M).

The simulations where both the A and B molecules contain 5 stickers (hereafter “A5:B5”) are shown in Fig. 7A while the A8:B8 simulations are shown in Fig. 7C. In both cases the phase boundary curves substantially away from the 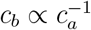 power law (black lines) expected for a strict *s* = 1 stoichiometry. However, the phase boundaries can be adequately fit by Eq. 34 using a two-step process in which *f*_*A*_ is first adjusted such that 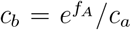 (black line) is tangent to the observed phase boundary. The second step is to adjust *α*_*A*_ to match the observed curvature (blue lines). Because the most reliable data for the phase boundary lie at the upper edge of the distribution, all fits were done visually (not by least squares regression).

#### Phase boundary curvature predicts composition of the dense phase

Having obtained *f*_*A*_ and *α*_*A*_ from the phase boundary, we can predict the composition of the dense phase without additional parameters. Fig. 7B,D shows the B:A ratio for both the monomers and dense phase. The red lines show the predicted composition of the dense phase, obtained by plugging Eq. 34 into Eq. 37. This is in good agreement with the observed composition (red points) of clusters containing *n* + *m* ≥ 20. Similar results were obtained for size cutoffs of 40 and 60, albeit with fewer simulations having clusters meeting the size criteria.

For asymmetric simulations, where the A and B molecules have different valences, the situation is slightly more complicated because the optimal composition *s* of the dense phase is not known *a priori*. This creates uncertainty in which power law should be used as the reference tangent to the phase boundary. However, inspection of Fig. 4 reveals a second condition that can be used to obtain *s*. Specifically, the −*s*^−1^ power law must be tangent to the phase boundary at the point of balanced chemical potentials *s* ln *c*_*a*_ = ln *c*_*b*_ (see Eq. 33). This second requirement again allows for a two-step fitting procedure where *f*_*A*_ and *s* are determined first, such that the power law and large *α* _*A*_ lines share a common tangent to the phase boundary, and then *α* _*A*_ is adjusted to match the observed curvature.

Fig. 8 compares the theory to the simulations of A9:B7, A10:B6, and A12:B4 molecular pairs. Again, the parameters obtained from fitting the phase boundary do an excellent job of predicting the composition of the dense phase across the phase boundary. Interestingly, the *s* values are far below what would be expected from the valence ratio. While the A9:B7 system has a valence ratio of 1.29, the obtained value of *s* was 1.02; for A10:B6 we fit 1.06 instead of the expected 1.67; and for A12:B3 we fit 1.10 instead of the expected 3.0. Reassuringly, the stoichiometry values obtained from matching the tangent of the phase boundary are close to the large cluster compositions in the simulations (see Fig. S2). Of course, the valence ratio will only give the correct stoichiometry in the strong-binding limit, where nearly all bonds are satisfied in the dense phase. In contrast, our simulations used a relatively weak affinity of 350 *µ*M where even the minority molecule only forms about half of its possible bonds (see Fig. S3). Under such conditions other contributions to the free energy, primarily the mixing entropy, will bias the dense phase composition toward more balanced ratios. The low values we found for *s* are most likely due to large activity coefficients in the dense phase that make it difficult to achieve the high concentrations needed for the low-valence species.

**Figure 8.**
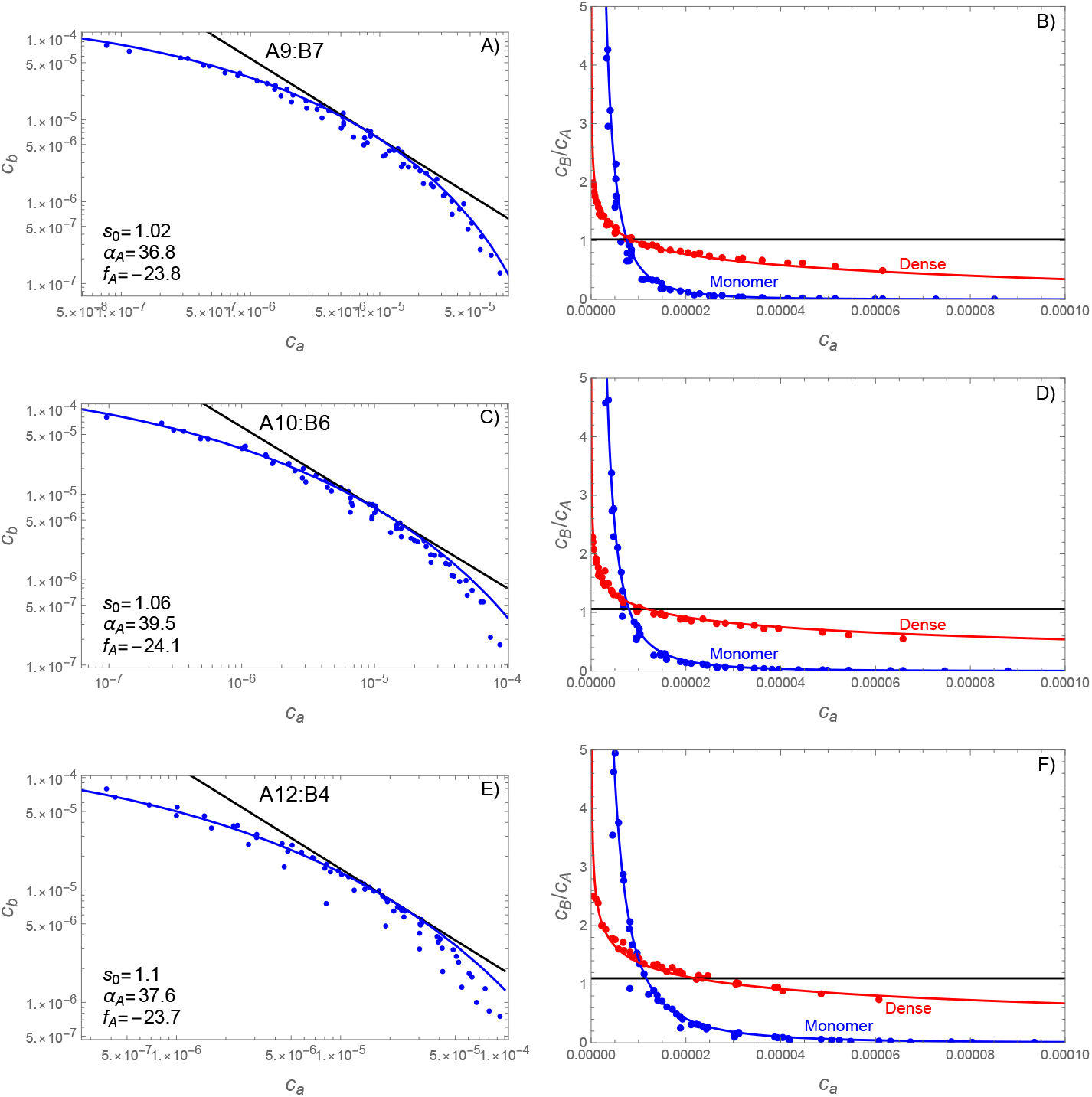
SpringSaLaD simulations reveal a curved phase boundary that predicts composition variations in the dense phase. Panels A,B show *v*_*A*_ = 9, *v*_*B*_ = 7; panels C,D show *v*_*A*_ = 10, *v*_*B*_ = 6; panels E,F show *v*_*A*_ = 12, *v*_*B*_ = 4;. Left panels (A,C,E) show the average concentration of free monomers for each of the 64 simulated mixing ratios. Fitting Eq. 34 to the uppermost points yields values for *f*_*A*_ and *α*_*A*_. Right panels (B,D,F) show the stoichiometric ratios B:A for both the monomers (blue) and dense phase (red). Note that the monomer lines and data (blue) are equivalent between panels with only a change in the vertical axis. The monomer ratio is calculated from Eq. 34 divided by *c*_*a*_ and the dense phase ratio is calculated from Eq. 37 with Eq. 34 substituted for *c*_*b*_. The stoichiometric composition of the dense phase agrees well with the composition of clusters *n* + *m* ≥ 20 molecules observed in the simulations. In all panels simulation data are shown as points, theory as a solid line, and concentrations are given in the dimensionless units *c*_*i*_ = *C*_*i*_/(1 M).

Inspection of Fig. 8F shows that the red, blue, and black lines do not intersect at a common point. As a result, when the monomer phase has the ideal stoichiometry for the dense phase *s* = 1.1, the dense phase has a composition ratio of ∼ 1.4. This is because the dense phase reaches the optimal composition with the chemical potential balance condition 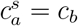 (see discussion following Eq. 37) and not when the monomer phase has the matching composition.

In Figs. 8C,E we see that the agreement between the observed phase boundary and the fitted line degrades with increasing distance from where the strict stoichiometry line (black) is tangent to the fitted curve. This disagreement is asymmetric with the theory falling below and above the observations on the left and right sides of the plot, respectively. This asymmetry suggests that the discrepancy between the fit and the theory may be due to anharmonic (*O* (*δ*^3^)) free energy contributions that were omitted in the approximation of Eq. 25. If this interpretation is correct, we would expect that the A-rich regime, where the two-phase region is more expanded, would be more permissive to variations in the dense phase stoichiometry than the B-rich side. Unfortunately, the simulations in the poorly fit regions did not have clusters above the *n* + *m* ≥ 20 size cutoff, which limits our ability to assess this explanation. Lowering the size cutoff to 10 (Fig. S5) suggests that the A-rich clusters do, in fact, have a greater propensity for stoichiometric variation. However, the inclusion of clusters with significant surface effects (i.e., the small cluster peak in Fig. 6A) lowers confidence in this observation.

### Dilute phase oligomers can be subtracted from dilute phase concentration measurements

In our simulation analysis, we circumvented the complication due to oligomers by directly measuring the concentration of monomers. However, this level of detail is not usually available from experiments. An alternative approach is to characterize the binding affinities in the dilute phase, and use the resulting equilibrium constants to compute the monomer concentrations.

We illustrate this approach using condensates formed from polyubiquitin and UBQLN2. ^38,42^ Fig. 9A shows dilute and dense phase concentrations measured for UBQLN2 and four variants of tetra-ubiquitin (Ub_4_) that differ in the linkage connecting the ubiquitin domains.^42^ Since each ubiquitin module has a single binding site for UBQLN2, the Ub_4_ binding polynomial is

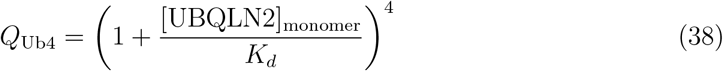

where [UBQLN2]_monomer_ is the concentration of UBQLN2 that is not bound to a ubiquitin module and *K*_*d*_ = 2.6 *µ*M is the mono-ubiquitin/UBQLN2 binding affinity.^38^ In using Eq. 38 to model the ensemble of solution oligomers, we are neglecting oligomers with two or more UB_4_ subunits or *>* 4 UBQLN2 molecules. Such approximations are valid when there is a hierarchy of interaction affinities^43^ so that it is expected that only the stronger interactions will contribute in the dilute phase. Another case worth of discussion is when the valence of a molecule is determined by homo-oligomerization.^24,44^ While a strict application of our theory would mandate that the unbound subunit takes the role of the monomer, if the homo-oligomers are sufficiently strong and monodisperse, it may be appropriate to neglect the small population of free subunits and treat the homo-oligomer as the monomeric building block.^24^

**Figure 9.**
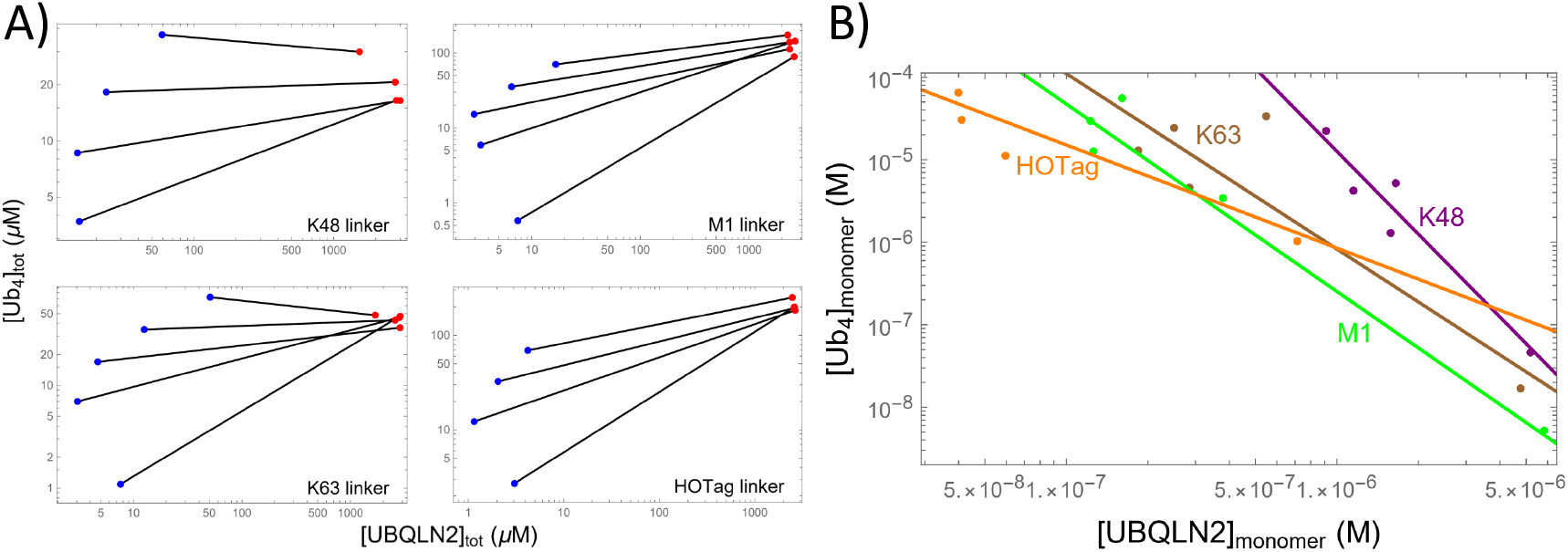
(A) UBQLN2/polyubiquitin phase diagrams, ^42^ plotted as a function of total concentrations, show re-entrant behavior as a function of Ub_4_ concentration. The four panels differ in the linkages used to connect the four ubiquitin modules. The non-physical crossing of tie lines is attributed to experimental uncertainty. (B) Subtracting the oligomer concentrations from the dilute phase (using Eqs. 39 and 43) reveal monomer concentrations consistent with the power law behavior expected from the solubiltiy product. K48: *f*_*A*_ = −17.2, *s* = 0.30; K63: *f*_*A*_ = −20.4, *s* = 0.47; M1: *f*_*A*_ = −20.5, *s* = 0.44; HOTag: *f*_*A*_ = −25.0, *s* = 0.80

From Eq. 38 the concentration of Ub_4_ that is unbound is

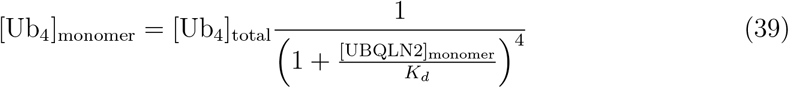

where [Ub_4_]_total_ is the total concentration of dilute phase Ub_4_. The total dilute phase concentration of UBQLN2 is the sum of the monomers and UBQLN2 bound in oligomers

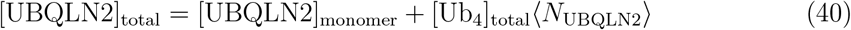

where ⟨ *N*_UBQLN2_ ⟩ is the average number of bound UBQLN2 molecules per Ub_4_, which is computed by

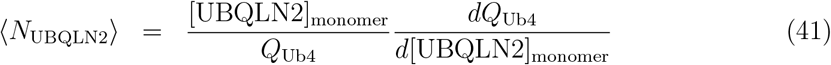

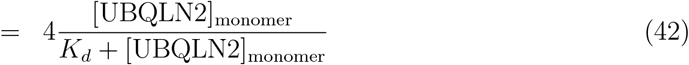

Inserting Eq. 42 into Eq. 40 yields a quadratic equation for UBQLN2 with the solution

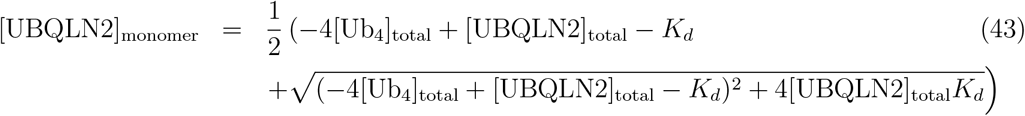

Eq. 43 can be inserted into Eq. 39 to obtain the concentration of monomer Ub_4_.

Applying Eqs. 39 and 43 to the dilute phase data in Fig. 9A reveals monomer concentrations that, when plotted on double logarithmic axes (Fig. 9B), are clustered along linear trend lines. Consistent with this linearity, measurements of the dense phase molecular composition show nearly constant stoichiometry over the range of study (Fig. S6). However, there is a significant discrepancy between the measured stoichiometry (0.01 to 0.1) and the fitted *s* values (0.3 to 0.8). Both of these ranges differ from the 0.25 value expected from the 1:4 valence ratio. There are several factors contributing to this. First, due to the clustering of points at low UBQLN2 concentration, the fitted slope is dominated by a single data point at high UBQLN2 concentration, which limits the statistical robustness of the fit. Furthermore, the HOTag variant lacks the highest concentration data point (Table S1), which further increases the uncertainty and likely explains why the HOTag slope is nearly double the other variants.

On the experimental side, concentration measurements in the dense phase come with large uncertainty due to the small volumes, high concentrations, and limited dynamic range of fluorescence detectors. For instance, fluorescence detection in three-dimensions is strongly affected by the point-spread-function,^45^ which is an important consideration for many cellular condensates that lie at or below the resolution limit of light microscopes. In these cases, accurate fluorescence measurements can be obtained by applying the point-spread-function but volume measurements are not feasible due to the subresolution size of these condensates. Previous work from our groups also found that accurate measurements of fluorescence intensity that fell within the linear range of the microscope detectors required unique imaging settings for each phase,^19^ supporting previous work that alluded to this technical limitation for accurately measuring partition coefficients.^46^

Measurement of the tie line slope has proven to be a useful tool to understand the balance of attractive and repulsive interactions driving phase separation.^47^ However, the difficulties described above can make tie lines difficult to measure. Our theory provides an alternative method in which the composition of the dense phase can be extracted from less technically challenging measurements of the dilute phase.

## Discussion

Our calculation shows that the solubility product is a robust description for salts due to two criteria that are often inappropriate for multi-component biomolecular condensates. The first of these is the neglect of soluble species larger than monomers, which is valid for salts due to a large surface tension suppressing clusters smaller than the critical nucleus. In contrast, biomolecular condensates tend to have comparatively low surface tension^48^ and a high propensity to form clusters in the sub-saturated regime ^20,49^ due to multiple factors including stoichiometry matching,^26,35^ conformational states, ^50^ and hierarchical assembly involving multiple interactions.^43^ The complication due to oligomers can be alleviated by considering the solution to be a three-state equilibrium between monomers, the condensed state, and the oligomer ensemble (Fig. 10). The equilibrium between the monomer and condensed phases resembles the power law of the solubility product and the sharpness of the condensation transition allows for straightforward calculation of the oligomer equilibrium in the sub-saturated regime.

**Figure 10.**
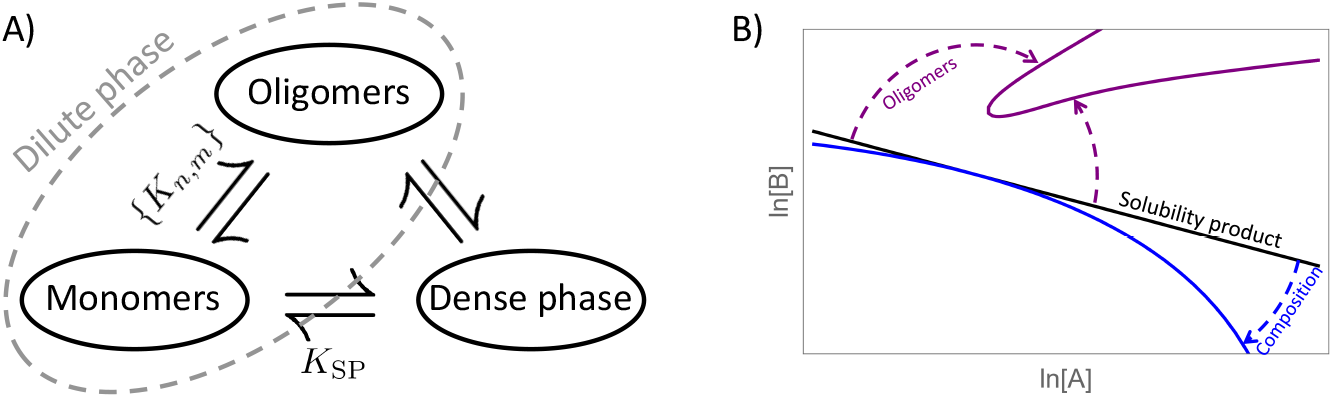
A) The equilibrium between monomers and oligomers can be calculated from dissociation constants while the equilibrium between monomers and the dense phase is described by the solubility product. The lack of a simple relationship between oligomers and the dense phase is responsible for the complexities associated with describing partitioning between the dense and dilute states. B) Schematic representation of the perturbations considered in this work. Oligomer formation and variation in the dense phase composition introduce opposite curvatures in the power law phase boundary that is expected from a simple solubility product.

**Figure 11.**
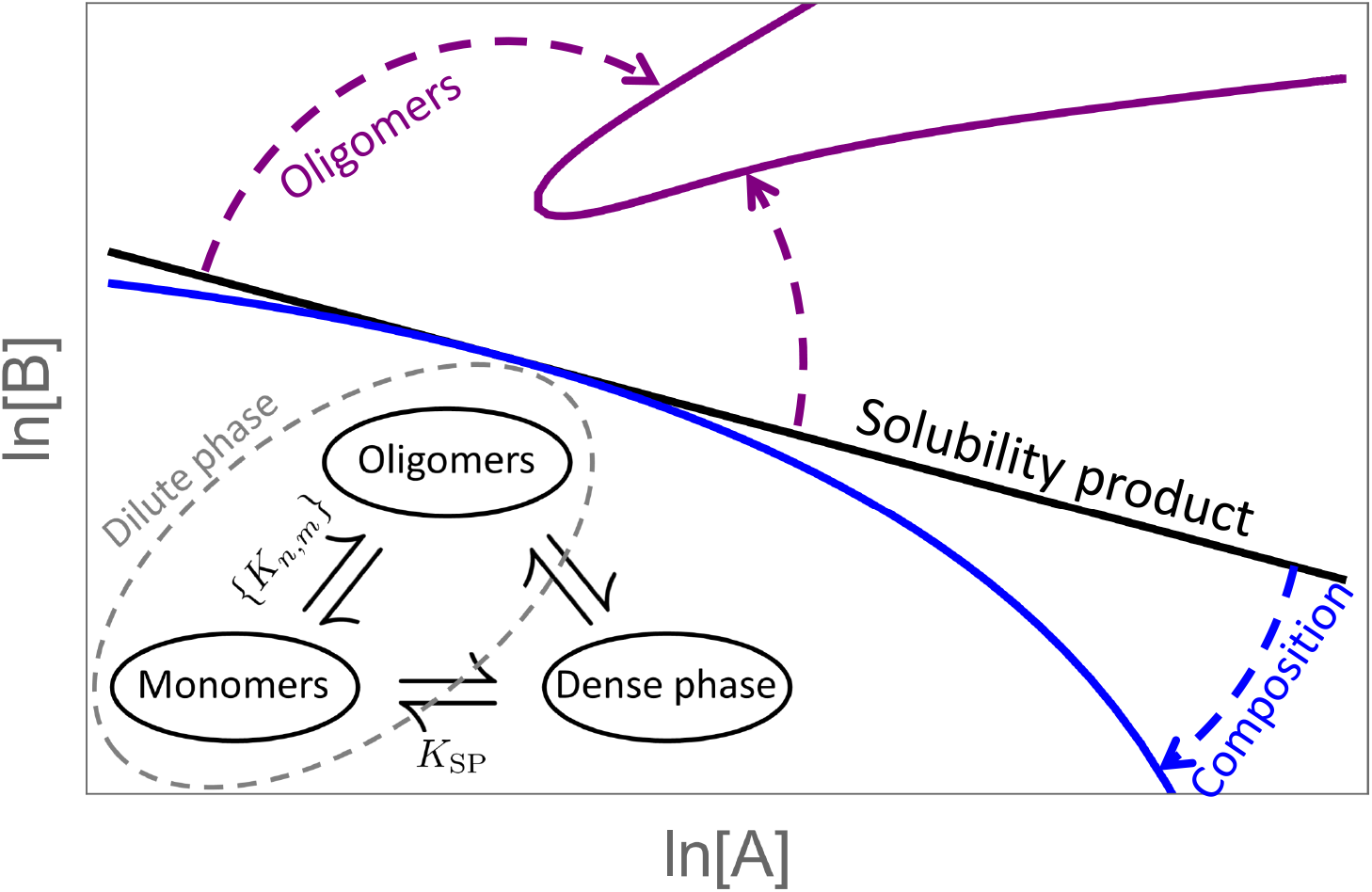
Table of Contents Figure

The second complication is that, unlike salts, biomolecular condensates do not have tightly constrained stoichiometry in the dense phase. Compositional variation lowers the free energy of the dense phase, expands the two-phase region of the phase diagram, and causes the monomer-condensate phase boundary to deviate from a simple power law. However, the power law deviation can be informative because it is predictive of how the composition of the dense phase varies. The solubility product relationship between monomers and the dense phase provides the second leg of the three state model shown in Fig. 10A. With two legs established, the full equilibrium is straightforward to calculate. The third leg, describing the oligomer/dense phase equilibrium, is less intuitive, but Eq. 21 can be used to compute the coexistence between an arbitrary oligomer and the dense phase.

Our formalism, based on the system partition function, differs from the more traditional approach, exemplified by Flory-Huggins, Cahn-Hilliard, or associative polymer theories, ^51–56^ of adopting a free energy that is a function of the local molecular concentrations. This difference brings advantages and disadvantages. For example, the explicit inclusion of small oligomers in our theory addresses the expected breakdown of mean-field approximations in the dilute phase; a difference that gives rise to our proposed three-state framework (Fig. 10A). Conversely, our formalism is not readily adapted to kinetics because the lack of spatial coordinates makes it difficult to account for mass transport. Similarly, because our theory does not specify the concentration of molecules within clusters, tie lines cannot be directly computed within our theory. However, a strength of analytic approaches is that shortcomings can be addressed by mixing models of different resolutions, such as the inclusion of oligomers in mean field models.^28^ In this regard, we expect our formalism to be particular flexible in that the cluster free energies *F* (*n, m*) are readily adapted to either specific molecular binding states, as in Eq. 38, or mean field treatments for large clusters. We envision that such a treatment could add the spatial degrees of freedom needed to compute tie lines.

The flexibility of the *F* (*n, m*) parameters allows our formalism to be applied to a wide range of systems ranging from salts to biomolecular condensates. Salts fall within the tight-binding limit where the free energies are dominated by binding energy, the interfacial tension is very high, and stoichiometric variation is strongly penalized (large *α*). In contrast, biomolecular condensates tend to have weaker binding energies, multiple mechanisms of binding, low surface tension, condensation driving forces dominated by entropy, ^24,33,57^ and are permissive of stoichiometric variations (small *α*). Generally, we expect factors like weak binding, high valence, conformational flexibility, and promiscuous interactions will contribute to greater stoichiometric variation.

It has been proposed that phase separating systems with two macromolecule components can be classified into four types. ^37^ Of these, our formalism is most readily applied to “cross-interaction-driven” phase separation. In cross-interaction systems, like our simulations, heterotypic interactions between the two molecule types are required to form the dense phase. Our theory is also readily applied to “scaffold-client” systems, such as polyubiquitin-UBQLN2 condensates. In such systems, the scaffold molecule is capable of condensing via homotypic interactions, which can introduce errors in the convergence of our composition integral. The concern is that, if the homotypic condensate is low enough in free energy, the *δ* integration may capture non-physical contributions from negative *n* or *m* states. Such states bring *attractive* translational entropy to the dense phase, leading to erroneous phase boundaries. In practice, however, the valence amplification enabled by the client/hub molecule tends to exponentially suppress the phase boundary of the heterotypic condensate below that of the homotypic condensate,^24,38^ which allows the composition integral to converge within the positive (*n, m*) region. The concern about the *δ* integral convergence increases with “cooperative” phase separation systems, in which both molecules are capable of phase separating independently. This makes it likely that the dense phase free energy minimum is too shallow (*α* is too small) for the composition integral to converge, especially in cases where the cooperative molecules are very similar. Finally, our theory is not applicable for “exclusive” phase separation, in which repulsive interactions between the macromolecules drive phase separation. Such systems are better handled by Flory-Huggins or Cahn-Hilliard theories, unless the repulsion can be absorbed into effective *F* (*n, m*) parameters.

In recent years, the search for phase separated compartments in cells has revealed a growing list of molecules that form puncta within cells. Since many of these puncta are quite small (< 100 nm), it raises the question of whether these structures should be considered biomolecular condensates, or simply clusters of molecules.^58^ Our decomposition of the solution into monomer, oligomer, and dense phases provides a framework to resolve this issue. Specifically, molecular assemblies should be considered condensates when they meet the linear scaling requirements necessary for inclusion in *Q*_den_. Assemblies that do not meet this criteria belong in *Q*_oligo_ and should be considered molecular clusters. There are at least two mechanisms by which an assembly might fail the criteria for *Q*_den_. First, generically we expect a surface contribution to the free energy that scales as *n*^2/3^, making it significant for small clusters but negligible for macroscopic phases. Second, biomolecular condensates often assemble from a variety of interaction motifs ranging from specific interactions between globular domains to weak interactions between IDRs. This will naturally lead to hierarchical assembly^43^ in which the stronger interactions form multimer clusters and the weaker interactions drive coalescence of multimer subunits. The relatively high affinity interactions responsible for the initial clusters are not representative of the free energy of the larger assembly and, therefore, these clusters should be included in *Q*_oligo_.

While our calculation provides intuition into factors that shape biomolecule phase boundaries, it does not account for many of the complexities of membraneless organelles in cells. Most severely, our calculation assumes the system is in equilibrium and, therefore, cannot account for the myriad of non-equilibrium and active processes that influence condensate formation. ^56^ We have also limited our calculation to two components. In contrast, biomolecular condensates can have hundreds of components that are enriched above the background level, although usually only a small number of scaffold species are necessary to reconstitute condensation *in vitro*.^59^ While it would be straightforward to generalize our method to more components, this would come at the cost of more parameters. Finally, many condensates have multiple structural states due to aging^60^ or biological responsiveness.^24,39,43^ Such transitions are due to multiple minima in the condensed phase free energy. Our calculation assumes a single free energy minimum and, therefore, cannot capture changes in the phase boundary due to a transition between conformational states. This deficiency could be exploited, however, because deviations from our theory could indicate the presence of biologically relevant states. For example, a change in the conformational state ^39^ would result in distinct sets of *f*_*A*_ and *α* parameters, and if the conformational change was induced by concentration,^24^ the curvature in the monomer phase boundary would show a kink at the point where the conformation changes.

## Conclusion

Many of the complexities in understanding the phase boundaries of biomolecular condensates can be attributed to the appearance of partitioning between two states, dense and dilute. It is conceptually easier, instead, to consider a three-state partitioning between monomers, the dense phase, and the oligomer ensemble (Fig. 10A).^19,28,34,61^ In the three-state picture, the monomer-oligomer equilibrium can be addressed with association constants, while the monomer-dense phase equilibrium is captured by the solubility product. In addition, we have shown that oligomers and dense phase stoichiometry distort the phase boundary in opposite ways; oligomers reduce the size of the two-phase region, while compositional variation expands it (Fig. 10B). The combination of the three-state model and the separation of oligomer and composition effects on the phase boundary allows for greater intuition into how molecular characteristics results in qualitatively different phase diagrams.

Supporting Information. Detailed calculations, simulation methods, and supplementary figures.

## Supporting information

Detailed derivations

## Acknowledgements

S.A.A., T.S., and J.D.S. acknowledge support from NIH grant R01GM141235. L.L. and A.C. acknowledge NIH grants R01GM132859 and R24GM137787.

J.A.D. acknowledges support from Natural Sciences and Engineering Research Council Discovery Grant RGPIN-2022-03274. J.D.S. would like to acknowledge helpful discussions with K. Stott and C. Castañeda.

Competing interests: Jonathon Ditlev is an advisor for Dewpoint Therapeutics and a scientific advisory board member for Neurophase Therapeutics.

