## Supplementary material for "Biomolecular phase boundaries are described by a solubility product that accounts for variable stoichiometry and soluble oligomers": Detailed derivations

#### Magic number effect

Consider a $(n, m)$ oligomer where $m/n$ is close to the stoichiometry of the dense phase. The oligomer and dense phase equilibrium conditions are

$$c_{n,m} = c_a^n c_b^m e^{-F(n,m)} \tag{S1}$$

$$c_a^n c_b^{ns} = e^{nf_A} \tag{S2}$$

where the second line is Eq. 8 raised to the $n$th power. Using $\epsilon = s - m/n$ as a small parameter, the phase boundary can be manipulated to find the concentration of the $(n, m)$ oligomer at the onset of phase separation

$$c_a^n c_b^m = e^{nf_A} c_b^{-n\epsilon} \tag{S3}$$

$$c_{n,m} e^{F(n,m)} = e^{nf_A} c_b^{-n\epsilon} \tag{S4}$$

Solving for the oligomer concentration we have

$$c_{n,m} = \exp[nf_A - F(n,m) - n\epsilon \ln c_b] \tag{S5}$$

We assume that the A valence is held fixed while the B valence is varied and that the solution mixing ratio is held fixed at a value $S = c_b/c_a$ as the valence is changed. In this case $c_a^n c_b^m = c_b^{n+m} S^{-n}$. Therefore the threshold oligomer concentration can be written as

$$c_{n,m} = \exp\left[nf_A - F(n,m) - \frac{n\epsilon}{n+m} \ln c_b^{n+m}\right] \tag{S6}$$

$$= \exp\left[nf_A - F(n,m) - \frac{n\epsilon}{n+m} \ln c_a^n c_b^m S^n\right] \tag{S7}$$

$$= \exp\left[nf_A - F(n,m) - \frac{n\epsilon}{n+m} \ln(c_{n,m} e^{F(n,m)} S^n)\right] \tag{S8}$$

This expression can be used recursively to obtain a power series in $\epsilon$. Applying the recursion iteratively gives

$$c_{n,m} = \exp\left[nf_A - F(n,m) - \frac{n\epsilon}{n+m}\ln(S^n e^{nf_A - \frac{n\epsilon}{n+m}\ln(c_{n,m}e^{F(n,m)}S^n)})\right] \tag{S9}$$

$$= \exp\left[nf_A - F(n,m) - \frac{n\epsilon}{n+m}\left(n\ln S + nf_A - \frac{n\epsilon}{n+m}\ln(c_{n,m}e^{F(n,m)}S^n)\right)\right] \tag{S10}$$

$$= \exp\left[nf_A - F(n,m) - \frac{n\epsilon}{n+m}(n\ln S + nf_A) + \left(\frac{n\epsilon}{n+m}\right)^2 \ln(c_{n,m}e^{F(n,m)}S^n)\right] \tag{S11}$$

$$= \exp\left[nf_A - F(n,m) - \frac{n\epsilon}{n+m}(n\ln S + nf_A) + \left(\frac{n\epsilon}{n+m}\right)^2 (n\ln S + nf_A) - \left(\frac{n\epsilon}{n+m}\right)^3 \ln(c_{n,m}e^{F(n,m)}S^n)\right] \tag{S12}$$

$$= \exp\left[nf_A - F(n,m) - (n\ln S + nf_A)\frac{n\epsilon}{n+m}\sum_{j=0}^{\infty}\left(\frac{-n\epsilon}{n+m}\right)^j\right] \tag{S13}$$

$$= \exp\left[nf_A - F(n,m) - (n\ln S + nf_A)\frac{n\epsilon}{n+m+n\epsilon}\right] \tag{S14}$$

$$= \exp\left[nf_A - F(n,m) - n(\ln S + f_A)\frac{s - m/n}{s+1}\right] \tag{S15}$$

$$= K(n,m)\exp\left[(n+m)\frac{f_A}{s+1} + \frac{m - ns}{s+1}\ln S\right] \tag{S16}$$

Eq. S16 allows for the calculation of the concentration of any oligomer along the phase boundary, provided $F(n,m)$ is known.

To understand Eq. S16, we note that the final term is zero when $c_a = c_b$. Defining $c_{ab}$ as the point on the phase boundary where $c_a = c_b$, we have $c_{ab}^{s+1} = e^{f_A}$ or $\ln c_{ab} = f_A/(s+_1)$. Therefore the leading term in Eq. S16 can be rewritten as $(n+m)\ln c_{ab}$, which is the translational entropy cost of forming the $(n,m)$ oligomer at this reference state. Therefore, the second term $(m - ns)(\ln S)/(s+1) = n\ln(c_a/c_{ab}) + m\ln(c_b/c_{ab})$ corrects the recruitment cost for deviations away from the reference state (this relationship follows from $c_{ab} = e^{f_A/(s+1)} = c_a^{1/(s+1)}c_b^{s/(s+1)}$).

In cases where the oligomer affinities are well characterized, Eq. S16 can be used to construct a parametric plot of the phase boundary as a function of the monomer ratio $S$. The location of the phase boundary at an arbitrary monomer ratio is

$$(c_A(S), c_B(S)) = \left( e^{\frac{f_A}{1+s}} S^{\frac{-s}{1+s}} + \sum^{\text{oligos}} n c_{n,m}(S), e^{\frac{f_A}{1+s}} S^{\frac{1}{1+s}} + \sum^{\text{oligos}} m c_{n,m}(S) \right) \tag{S17}$$

where the leading terms are the monomer concentrations expressed as a function of $S$.

We can generalize the magic number analysis for arbitrary oligomers. We start with the first order approximation for $c_{n,m}$ by taking Eq. S11 and dropping terms of order $\epsilon^2$.

$$c_{n,m} = \exp\left[ n f_A - F(n,m) - \frac{n^2 \epsilon}{n+m}(f_A + \ln S) \right] \tag{S18}$$

In the tight binding limit the oligomer free energy is dominated by the maximum number of bonds

$$F(n,m) = f_1 \min(v_A n, v_B m) \tag{S19}$$

$$= f_1 n v_A \min\left(1, \frac{m}{ns}\right) \tag{S20}$$

$$= f_1 n v_A \min\left(1, \frac{1}{1+\frac{n\epsilon}{m}}\right) \tag{S21}$$

where $f_1$ is the free energy of a single intermolecular bond. Therefore, for $\epsilon < 0$ the threshold oligomer concentration varies as

$$c_{n,m} = \exp\left[ n f_A - f_1 n v_A - \frac{n^2 \epsilon}{n+m}(f_A + \ln S) \right] \tag{S22}$$

which increases exponentially as $\epsilon$ approaches zero. For $\epsilon > 0$ we have $F(n,m) \simeq f_1 n v_A (1 - n\epsilon/m)$, so in the small $\epsilon$ limit the oligomer concentration at the phase boundary varies as

$$c_{n,m} = \exp\left[ n f_A - n f_1 v_A - \frac{n^2 \epsilon}{n+m}(f_A + \ln S) + \frac{n^2 \epsilon}{m} f_1 v_A \right] \tag{S23}$$

which *decreases* exponentially provided that $f_1 v_A < (f_A + \ln S)m/(n+m)$. This inequality can be interpreted as follows. The left side is the binding energy of the oligomer normalized by the number of A molecules. Similarly, $f_A$ is the free energy per A molecule in the dense phase. However, at equilibrium $f_A$ is balanced against the entropic cost of recruiting molecules to the condensate. Therefore, the right side represents the entropic cost of recruiting the B molecules to the dense phase. This interpretation is supported by the $m/(n+m)$ factor that selects only the B molecules in a stoichiometric unit and the $\ln S$ factor that corrects the translational entropy cost for the solution stoichiometry. Therefore, oligomer concentration has a magic number peak when the benefit of a B molecule binding in the oligomer state outweighs the translational entropy of recruitment to the dense phase.

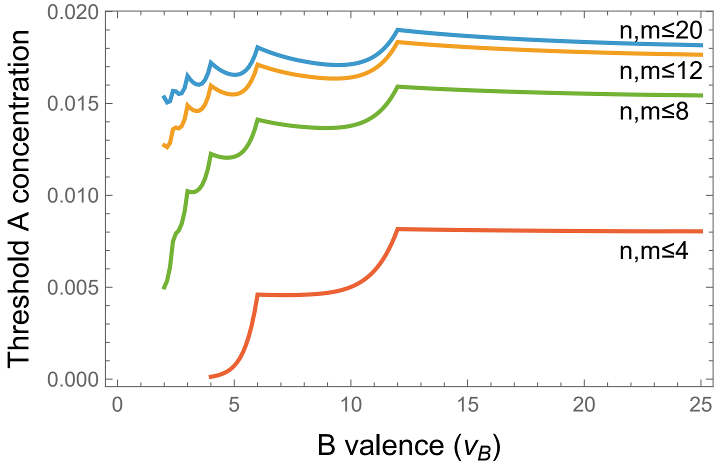


Figure S1: Adding A molecules of valence $v_A = 12$ to a fixed B concentration $C_B = 1$ M reveals magic number peaks at integer ratios of the molecular valence $v_A/v_B$. The different color lines show the effect of adding more oligomers to the model, with the maximum number of molecules in the oligomer shown on the graph (i.e., the $n, m \leq 20$ model includes $20 \times 20 = 400$ oligomer states in $Q_{\text{oligo}}$). Adding more oligomers to the model increases the threshold concentration because more A molecule is bound in oligomer states. The threshold concentration is obtained by numerically solving for the monomer concentrations that satisfy the predetermined total concentrations (via Eq. 11) and exceed the threshold condition $c_a c_b^s = e^{f_A}$. Curves are computed with the tight binding approximations $s = v_A/v_B$ and $F(n, m) = f_1 \min[n v_A, m v_B]$ with parameters $f_1 = -1$ and $f_A = -18$.

In agreement with this expectation, Fig. S1 shows magic number peaks for all integer ratios of $m/n$ as the valence of B is varied. The absence of peaks for oligomers with more than one A molecule can be attributed to holding the B concentration fixed ($C_B = 1$ M) as we varied $C_A$. This leads to very high $S$ values that strongly penalize the formation of oligomers with more than one A molecule.

### Evaluation of partition function integrals

Our goal is to evaluate the double integral

$$Q_{\text{den}} = \int_0^\infty \int_0^{\pi/2} c_a^{r\cos\theta} c_b^{r\sin\theta} e^{-F(r,\theta)} r dr d\theta \tag{S24}$$

The chemical potential terms can be Taylor expanded around $\theta_0$ to obtain an expression consistent with the second-order expansion of the free energy. For the A molecules we have

$$\begin{aligned} n &= r\cos\theta & \text{(S25)} \\ &= r(\cos\delta\cos\theta_0 - \sin\delta\sin\theta_0) & \text{(S26)} \\ &= \frac{r}{\sqrt{1+s^2}}\left(1 - s\delta - \frac{\delta^2}{2}\right) & \text{(S27)} \end{aligned}$$

and for the B molecules

$$\begin{aligned} m &= r\sin\theta & \text{(S28)} \\ &= r(\sin\delta\cos\theta_0 + \cos\delta\sin\theta_0) & \text{(S29)} \\ &= \frac{r}{\sqrt{1+s^2}}\left(s + \delta - s\frac{\delta^2}{2}\right) & \text{(S30)} \end{aligned}$$

where the last steps used the relations

$$\begin{aligned} \cos\theta_0 &= \frac{1}{\sqrt{1+s^2}} & \text{(S31)} \\ \sin\theta_0 &= \frac{s}{\sqrt{1+s^2}} & \text{(S32)} \end{aligned}$$

The expanded free energy (Eq. 24) and chemical potentials Eqs. S27 and S30 can be inserted back into the expression for the partition function

$$Q_{\text{den}} = \int_0^\infty \int_0^{\pi/2} c_a^{r\cos\theta} c_b^{r\sin\theta} e^{-F(r,\theta)} r dr d\theta \tag{S33}$$

$$\simeq \int_0^\infty \int_0^{\pi/2} e^{-r\delta^2 a + r\delta b - rc} r dr d\delta \tag{S34}$$

where

$$a = \frac{1}{2\sqrt{1+s^2}} \left[\alpha\sqrt{1+s^2} + \ln c_a + s \ln c_b\right] \tag{S35}$$

$$b = \frac{1}{\sqrt{1+s^2}} \left(\ln c_b - s \ln c_a\right) \tag{S36}$$

$$c = \frac{1}{\sqrt{1+s^2}} \left(f_A - \ln c_a - s \ln c_b\right) \tag{S37}$$

Next we complete the square in the exponent

$$Q_{\text{den}} = \int_0^\infty \int_0^{\pi/2} e^{-r\left(\delta^2 a - \delta b + c\right)} r dr d\delta \tag{S38}$$

$$= \int_0^\infty \int_0^{\pi/2} e^{-r\left(\delta\sqrt{a} - \frac{b}{2\sqrt{a}}\right)^2 + r\frac{b^2}{4a} - rc} r dr d\delta \tag{S39}$$

$$= \int_0^\infty e^{r\frac{b^2}{4a} - rc} \int_0^{\pi/2} e^{-ra\left(\delta - \frac{b}{2a}\right)^2} r dr d\delta \tag{S40}$$

To perform the integral over $\delta$ we extend the limits of integration to $\pm\infty$ so that the integral yields $(\pi/ra)^{1/2}$. While the extended limits introduce error by admitting regions where $n, m$ are negative, contributions from these regions will be strongly suppressed by the unfavorable free energies at extreme stoichiometries. Therefore, this error is likely smaller than the neglect of anharmonic terms.

Now we perform the integral over $r$

$$Q_{\mathrm{den}} = \int_0^\infty \sqrt{\frac{\pi}{ra}} e^{r\frac{b^2}{4a} - rc} r dr \tag{S41}$$

$$= \sqrt{\frac{\pi}{a}} \frac{1}{\left(c - \frac{b^2}{4a}\right)^{3/2}} \int_0^\infty \sqrt{x} e^{-x} dx \tag{S42}$$

$$= \frac{\pi}{2\sqrt{a}\left(c - \frac{b^2}{4a}\right)^{3/2}} \tag{S43}$$

where the integration variable is $x = r(c - \frac{b^2}{4a})$ and the final integral is $\Gamma(3/2) = \sqrt{\pi}/2$. Here we have introduced another approximation by extending the integration range to zero. This captures terms in the oligomer range, however, the divergence that determines the phase boundary comes from the large $r$ portion of the integral.

Inserting the values for $a$, $b$, and $c$ into Eq. S43, we have

$$Q_{\mathrm{den}} = \frac{\pi(1+s^2)}{\sqrt{2\left(\alpha_A + \ln\left(c_a c_b^s\right)\right)}\left(f_A - \ln\left(c_a c_b^s\right) - \frac{\left(\ln \frac{c_b}{c_a^s}\right)^2}{2\left(\alpha_A + \ln\left(c_a c_b^s\right)\right)}\right)^{3/2}} \tag{S44}$$

In this expression we see that the integral approximation to the partition function has introduced a second source of error in addition to extending the limits of the $\delta$ integration. Specifically, the above expression will not quantitatively reproduce the tight stoichiometry result. This can be seen by taking the limit $\alpha_a \to \infty$, which causes the partition function to vanish. This happens because the Gaussian that results from Taylor expanding the free energy $F(r, \theta) \simeq r(f + \alpha\delta^2)$ has vanishing width, and hence vanishing area, in the limit $\alpha \to \infty$. The denominator in Eq. S44 is somewhat different than the tight stoichiometry limit Eq. 7, although this is partly due to the integral approximation of the summation. Evaluating Eq. 7 using an integral, rather than a sum, yields a denominator $f_A - \ln c_a c_b^s$, which is equivalent to the small argument expansion of the exponential. Another significant change is the increase in the exponent of the divergent term from $-1$ to $-3/2$, which comes from including the second dimension in the partition sum.

### Phase boundary

We can use the solubility relation (Eq. 33) to derive a condition for the phase boundary.

$$c_a c_b^s = \exp\left[f_A - \frac{1}{2\alpha'}\left(\ln\frac{c_b}{c_a^s}\right)^2\right] \tag{S45}$$

With the definitions $x = \ln c_a$ and $y = \ln c_b$ this becomes

$$e^x e^{sy} = \exp\left[f_A - \frac{1}{2\alpha'}\left(y - sx\right)^2\right] \tag{S46}$$

Taking the logarithm of both sides and rearranging

$$2\alpha'(x + sy - f_A) \quad = \quad -\left(y - sx\right)^2 \tag{S47}$$

which is a quadratic equation for $y$. The physically relevant root is

$$y \quad = \quad sx - \alpha' s + \sqrt{(sx - \alpha' s)^2 - 2\alpha'(x - f_A) - s^2 x^2} \tag{S48}$$

$$\quad = \quad s\alpha'\left(-1 + \frac{x}{\alpha'} + \sqrt{1 - \frac{2x}{\alpha'} + \frac{2f_A}{\alpha' s^2} - \frac{2x}{\alpha' s^2}}\right) \tag{S49}$$

In the limit of large $\alpha'$ we recover the slope of $-s^{-1}$ expected for a power law

$$y \simeq \frac{2f_A}{s} - \frac{x}{s} \tag{S50}$$

Replacing $x$ and $y$ in Eq. S49 yields the phase boundary in the main text (Eq. 34).

It is straightforward, albeit slightly more cumbersome, to derive the phase boundary without employing the recursive approximation $\alpha' \simeq \alpha_A + \ln c_a c_b^s$. However, this more complicated expression yielded similar results in our comparison with simulations.

### Dense phase stoichiometry

To compute the stoichiometric composition of the dense phase, we begin with the dense phase partition function (Eq. S44). The total number of molecules in the dense phase can be computed using

$$c_A^{\text{den}} = c_a \frac{dQ_{\text{den}}}{dc_a} \tag{S51}$$

$$c_B^{\text{den}} = c_b \frac{dQ_{\text{den}}}{dc_b} \tag{S52}$$

We note that the partition function has the form

$$Q_{\text{den}} = \frac{\pi(1+s^2)}{g^{1/2}h^{3/2}} \tag{S53}$$

so that taking a derivative gives

$$Q'_{\text{den}} = Q_{\text{den}} \left( -\frac{g'}{2g} - \frac{3h'}{2h} \right) \tag{S54}$$

We are interested in the composition of the dense phase, which will dominate the solution when $h \simeq 0$. Under these conditions the derivative is dominated by the second term of Eq. S54. The stoichiometry of the dense phase is therefore

$$\frac{c_B^{\text{den}}}{c_A^{\text{den}}} = \frac{c_b \frac{dQ}{dc_b}}{c_a \frac{dQ}{dc_a}} \simeq \frac{c_b \frac{dh}{dc_b}}{c_a \frac{dh}{dc_a}} \tag{S55}$$

Evaluating the derivatives gives

$$c_a \frac{dh}{dc_a} = c_a \left( -\frac{1}{c_a} - \frac{-s \frac{1}{c_a} \ln \frac{c_b}{c_a^s}}{\alpha_A + \ln(c_a c_b^s)} + \frac{\frac{1}{c_a} \left( \ln \frac{c_b}{c_a^s} \right)^2}{2 \left( \alpha_A + \ln\left(c_a c_b^s\right) \right)^2} \right) \tag{S56}$$

$$\simeq \left( -1 - \frac{-s \ln \frac{c_b}{c_a^s}}{\alpha'} + \frac{\left( \ln \frac{c_b}{c_a^s} \right)^2}{2\alpha'^2} \right) \tag{S57}$$

$$c_b \frac{dh}{dc_b} = c_b \left( -\frac{s}{c_b} - \frac{\frac{1}{c_b} \ln \frac{c_b}{c_a^s}}{\alpha_A + \ln(c_a c_b^s)} + \frac{\frac{s}{c_b} \left( \ln \frac{c_b}{c_a^s} \right)^2}{2 \left( \alpha_A + \ln(c_a c_b^s) \right)^2} \right) \tag{S58}$$

$$\simeq \left( -s - \frac{\ln \frac{c_b}{c_a^s}}{\alpha'} + \frac{s \left( \ln \frac{c_b}{c_a^s} \right)^2}{2\alpha'^2} \right) \tag{S59}$$

The quantity $\ln(c_b c_a^{-s})/\alpha'$ is the driving force for composition fluctuations divided by the fluctuation stiffness. This can be used as a small parameter in the weak perturbation limit, implying that the third terms in the above equations are negligible. Keeping only the first two terms, we find that the the stoichiometric ratio is

$$\frac{c_B^{\mathrm{den}}}{c_A^{\mathrm{den}}} = \frac{s + \frac{1}{\alpha'} \ln \frac{c_b}{c_a^s}}{1 - \frac{s}{\alpha'} \ln \frac{c_b}{c_a^s}} \tag{S60}$$

$$\simeq s + \frac{(1+s)}{\alpha'} \ln \frac{c_b}{c_a^s} \tag{S61}$$

### SpringSaLaD Simulations

SpringSaLaD software[1] has been used in multiple publications[2–6] to study clustering of multivalent biomolecules and its implications in phase separation. In this framework, each molecule is represented by a collection of beads connected by harmonic bonds (springs). Motion of particles are governed by an overdamped Langevin equation. The springs transmit force to the connected beads; additionally, a random force (uncorrelated in time) is applied to each bead at each timestep whose magnitude is determined by the diffusion coefficient assigned to that bead. Each bead has a radius and the simulations are conducted in a cubic box with reflective boundaries.

In SpringSaLaD, certain beads engage in binding interactions while others serve as structural sites to mimic the actual molecular geometry. Each binding interaction is assigned association and dissociation rates whose ratio determines the macroscopic equilibrium constant. These rates are then converted to microscopic probabilities[1] in a thermodynamically consistent manner which guarantees the equilibrium distribution of molecular species. When particles diffuse in the simulation volume, reversible binding interactions are modelled with a Smoluchowski-like framework. When complementary binding sites are within a cutoff radius, they form a bond based on the probability computed from the association rate, particle radius, and diffusion coefficient. Similarly, the bond is broken with the probability calculated from the dissociation rate. Our model is not intended to replicate a specific system, however, the parameters are motivated by previous studies.[2,7]

In our simulation setup, we first consider a pair of octavalent molecules (A8, B8, Fig. 1A). Each molecular type has 8 stickers (binding sites) and 24 spacers (structural sites). The binding affinity between sticker pairs is 350 $\mu$M (association rate = 10 $\mu M^{-1}s^{-1}$ and dissociation rate = 3500/s). We used a bead radius = 1 nm, diffusion coefficient = 2 $\mu m^2/s$, and the length between two consecutive beads is 2.5 nm. Additional details about the simulation are available at `https://github.com/achattaraj/softSP_theory/tree/main`.

We titrate up molecular count in a fixed volume to change the concentration. For all simulations except A5:B5 we used volume = $120 \times 120 \times 120 nm^3$ and the molecular counts, $N_A$ and $N_B$ are taken from {15, 30, 45, 60, 75, 90, 105, 120}. As a reference, in the given volume, 1 molecule $\sim$ 0.96 $\mu$M. We create a grid of concentrations (64 points) by taking all combinations of $N_A$ and $N_B$ from the aforementioned list. For the A5:B5 simulations the simulation volume is $100 \times 100 \times 100 nm^3$ and the molecular counts, $N_A$ and $N_B$ are taken from {20, 40, 60, 80, 100, 120, 140, 160, 180}.

To create asymmetric molecular pairs, we extended (or reduced) the molecular length

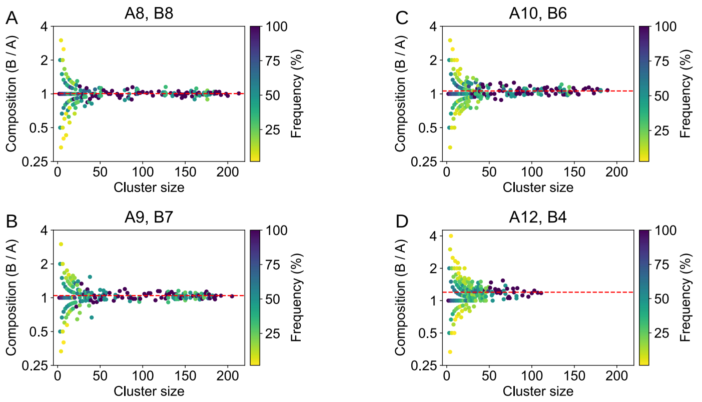


Figure S2: Summary of cluster composition for asymmetric (length and valence) molecule pairs in simulations with $N_A = N_B = 120$ molecules. The average composition for clusters larger than 50 molecules is given by the red dashed lines. The average values are 1.01 for A8:B8, 1.04 for A9:B7, 1.06 for A10:B6, and 1.20 for A12:B4.

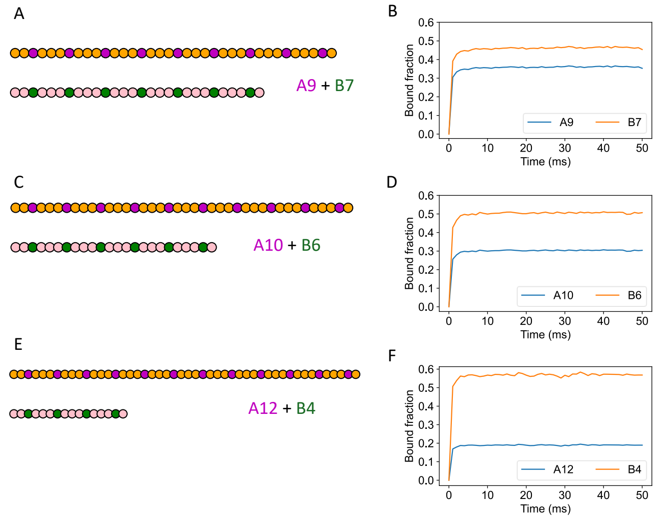


Figure S3: Panels A, C, and E show a schematic representation of the sticker and spacer patterns for the asymmetric simulations. Panels B, D, and F show bond saturation for the corresponding system. The bond saturation is $\sim 50\%$ for the minority component and even lower for the majority component. This low bond saturation allows for variations in the dense phase stoichiometry.

keeping the sticker:spacer ratio and total number of beads fixed. The chain characteristics are summarized using the notation $L(i, j)$ where $i$ and $j$ are the sticker and spacer count per molecule ($L = i + j$) respectively (see Fig. S3).

A8 + B8: L(8,24) + L(8,24)

A9 + B7: L(9,27) + L(7,21)

A10 + B6: L(10,30) + L(6,18)

A12 + B4: L(12,36) + L(4,12)

As the clusters grow in size, their compositions converge to a fixed value (Fig. S2). Smaller clusters show significant variation, but large clusters tend to have similar compositions.

Due to the difference in valence (sticker stoichiometry), the asymmetric simulations show different degrees of bond saturation for A-type and B-type chains (Fig. S3B,D,F). This difference grows as the sticker stoichiometry gets more imbalanced.

To test the effect of finite system's size, we considered three sizes for a given system (A8 + B8 or A10 + B6). Keeping the total concentration unchanged (Fig. S4A), we computed the cluster size distributions (Figs. S4B,C) and monomer concentrations (Fig. S4D,E). Overall, the cluster distributions are insensitive to the system size, with average cluster compositions close to one another. For monomer concentrations, the distributions overlap significantly, although we notice a slight (note the expanded scale of the Y-axes) upward trend in the median free concentrations with larger box sizes. Most likely, the trend is a consequence of the reflective boundary conditions. Due to reflections at the boundary, simulation walls and the excluded volume of the molecules can slightly increase the "effective concentrations". With larger simulation volume, surface to volume ratio goes down which lowers this "edge-effect". As a result, the effective molecular concentration is slightly higher in a smaller box, shifting the binding equilibrium (free $\leftrightarrow$ bound) towards the bound states, resulting in lower free concentrations.

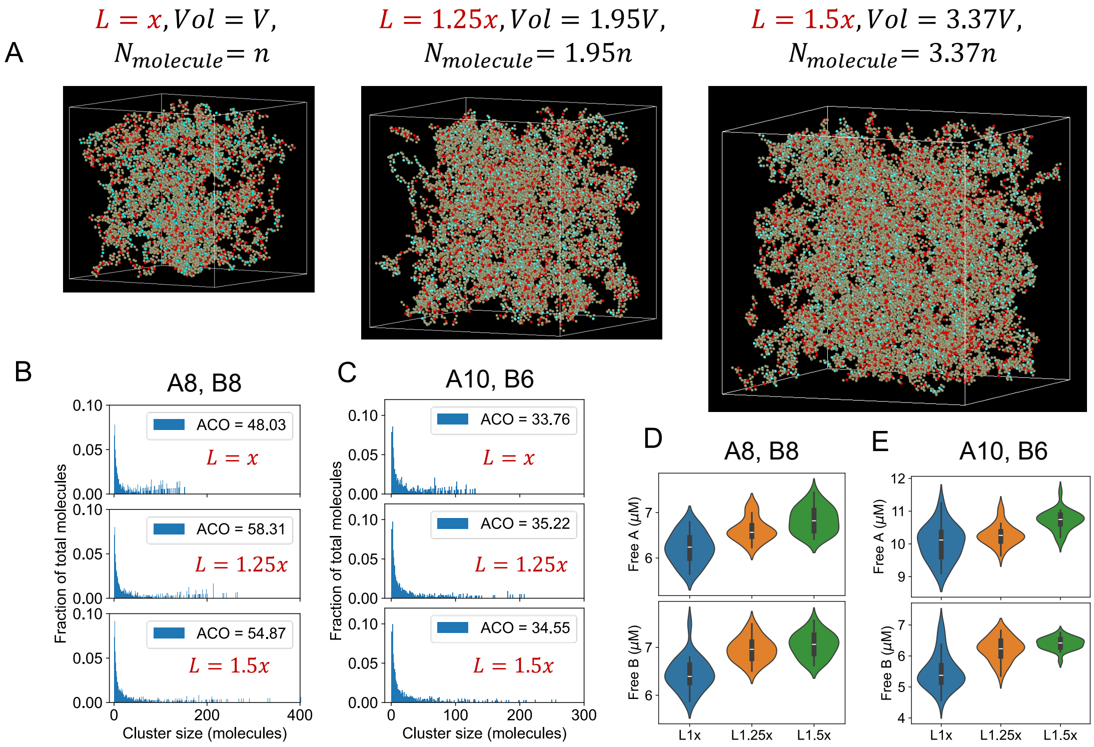


Figure S4: Summary of simulation finite size effects. (A) Three systems with different sizes are simulated. Size of the simulation volume $V$ and molecular counts ($N_{\text{molecule}}$) are increased proportionally to keep the concentration ($N_{\text{molecule}}/V$) fixed. $L$ is the length of the cubic box, $V = L^3$. For the reference system, $L = x = 120$nm, $N_{\text{molecule}} = N_A + N_B$, $N_A = N_B = 100$. These size titration simulations are done for two valencies: A8 + B8, and A10 + B6. (B,C) Cluster size distributions (ACO = average cluster occupancy), (D,E) Free molecular (monomer) concentrations, for two valencies at three system sizes. In D,E, standard violin plots are used to display the distribution. Violin: data density, white dot: median, bold bar in center: interquartile range (IQR, 25th–75th percentile), thin Black Line: Whiskers, extending 1.5 * IQR.

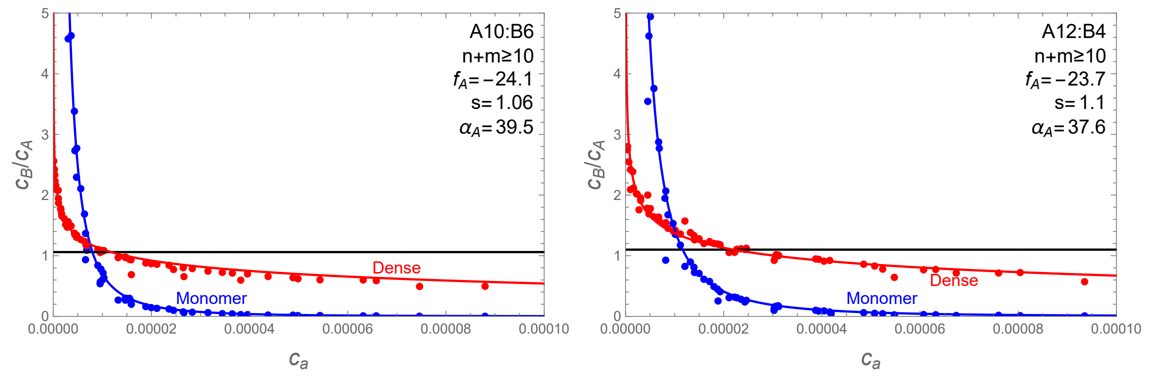


Figure S5: Composition plots for the A10:B6 and A12:B4 systems with a dense phase cutoff of $n + m \geq 10$ and the same parameters used in Fig. 8. The smaller cutoff (compared to $n+m \geq 20$ used in the main text) results in more simulations meeting the size threshold. The systematic deviation of the simulations below the red theory line for large $c_a$ is suggestive of a $\delta^3$ contribution to the cluster partition function.

### UBQLN2/polubiquitin

Parameter fitting was done with the LinearModelFit algorithm in Wolfram Mathematica. The obtained values are: K48: $f_A/s = -57.6 \pm 7.3$, $s^{-1} = 3.35 \pm 0.55$; K63: $f_A/s = -43.3 \pm 9.0$, $s^{-1} = 2.12 \pm 0.62$; M1: $f_A/s = -46.1 \pm 3.9$, $s^{-1} = 2.25 \pm 0.26$; HOTag: $f_A/s = -31.6 \pm 4.3$, $s^{-1} = 1.25 \pm 0.27$. Reported standard errors do not account for the propagation of experimental uncertainty.

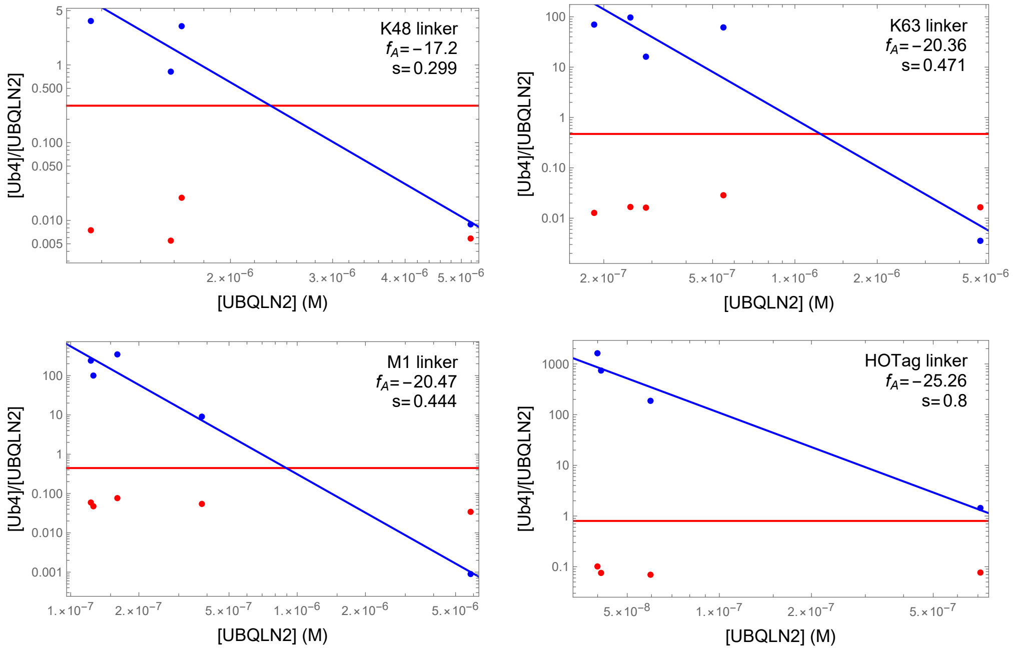


Figure S6: Composition ratios for the dilute phase monomers (blue) and dense phase (red) for each of the $Ub_4$ linker variants. Parameters for the blue and red lines are obtained from the fits in Fig. 9B. Blue points are calculated from Eqs. 39 and 43 (see Table). Dense phase stoichiometries (red points) are from [8] and the red lines are the predicted stoichiometries from the Fig. 8B fits. Both the red points and red lines differ from the expected stoichiometric ratio of 0.25.

| linker | Ub : UBQLN2 | $[\mathrm{UBQLN2}]_{\mathrm{dil}}$ | $[\mathrm{Ub_4}]_{\mathrm{dil}}$ | $[\mathrm{UBQLN2}]_{\mathrm{mono}}$ | $[\mathrm{Ub_4}]_{\mathrm{mono}}$ | $[\mathrm{UBQLN2}]_{\mathrm{den}}$ | $[\mathrm{Ub_4}]_{\mathrm{den}}$ |
|---|---|---|---|---|---|---|---|
| K48 | 0. | $19.62 \pm 0.28$ | | | | $2809 \pm 99$ | |
| | 0.2 | $15.07 \pm 0.20$ | $3.71 \pm 0.72$ | 5.187 | 0.05 | $2817 \pm 19$ | $65.94 \pm 9.20$ |
| | 0.5 | $14.63 \pm 0.17$ | $8.63 \pm 0.27$ | 1.579 | 1.294 | $2996. \pm 138$ | $65.8 \pm 25.7$ |
| | 1. | $23.6 \pm 1.5$ | $18.27 \pm 0.98$ | 1.150 | 4.224 | $2764 \pm 132$ | $83 \pm 22$ |
| | 2. | $59.1 \pm 3.5$ | $37.0 \pm 2.6$ | 1.648 | 5.196 | $1532 \pm 286$ | $120 \pm 52$ |
| | 4. | $78.06 \pm 3.83$ | $74.27 \pm 5.09$ | 0.9120 | 22.31 | | |
| K63 | 0. | $17.56 \pm 1.76$ | | | | $2559 \pm 169$ | |
| | 0.2 | $7.590 \pm 0.5698$ | $1.091 \pm 0.985$ | 4.767 | 0.017 | $2873 \pm 368$ | $189.2 \pm 49.4$ |
| | 0.5 | $3.041 \pm 0.4963$ | $6.968 \pm 0.741$ | 0.2852 | 4.595 | $2833 \pm 228$ | $183.5 \pm 42.6$ |
| | 1. | $4.679 \pm 0.427$ | $16.97 \pm 0.70$ | 0.1844 | 12.90 | $2863 \pm 370$ | $145.8 \pm 67.1$ |
| | 2. | $12.55 \pm 1.82$ | $35.02 \pm 1.62$ | 0.2502 | 24.25 | $2580 \pm 122$ | $172.2 \pm 67.7$ |
| | 4. | $50.88 \pm 19.90$ | $72.31 \pm 1.79$ | 0.5477 | 33.66 | $1693 \pm 252$ | $193.3 \pm 74.1$ |
| M1 | 0. | $15.91 \pm 2.48$ | | | | $2481 \pm 129$ | |
| | 0.2 | $7.417 \pm 1.479$ | $0.575 \pm 0.552$ | 5.827 | 0.005211 | $2615 \pm 190$ | $357.3 \pm 121.3$ |
| | 0.5 | $3.389 \pm 0.593$ | $5.908 \pm 2.996$ | 0.3794 | 3.426 | $2667 \pm 53.35$ | $579.8 \pm 83.7$ |
| | 1. | $2.943 \pm 0.572$ | $15.24 \pm 0.95$ | 0.1260 | 12.61 | $2387 \pm 134$ | $451.2 \pm 90.5$ |
| | 2. | $6.516 \pm 0.730$ | $35.45 \pm 3.14$ | 0.1228 | 29.48 | $2380 \pm 192$ | $561.9 \pm 64.6$ |
| | 4. | $16.62 \pm 2.38$ | $70.7 \pm 8.1$ | 0.1607 | 55.62 | $2284 \pm 75$ | $697 \pm 163$ |
| HOTag | 0. | $16.47 \pm 1.89$ | | | | $2419 \pm 113$ | |
| | 0.2 | $8.811 \pm 2.028$ | | | | $2622 \pm 137$ | $466.3 \pm 54.1$ |
| | 0.5 | $3.045 \pm 0.568$ | $2.708 \pm 1.193$ | 0.7132 | 1.027 | $2597 \pm 73.64$ | $797.5 \pm 94.6$ |
| | 1. | $1.153 \pm 0.746$ | $12.21 \pm 2.16$ | 0.05956 | 11.15 | $2644 \pm 43$ | $734.9 \pm 129.4$ |
| | 2. | $2.054 \pm 0.709$ | $32.38 \pm 3.43$ | 0.04105 | 30.41 | $2567 \pm 195$ | $774.2 \pm 147.2$ |
| | 4. | $4.216 \pm 0.655$ | $69.00 \pm 9.07$ | 0.03995 | 64.92 | $2474 \pm 246$ | $1002 \pm 99$ |

Table S1: Measured concentration in the dilute and dense phases from Dao et al.[8] along with the monomer concentrations calculated from Eqs. 39 and 43.
